# Variable species densities are induced by volume exclusion interactions upon domain growth

**DOI:** 10.1101/061341

**Authors:** Robert J. H. Ross, C. A. Yates, R. E. Baker

**Affiliations:** Wolfson Centre for Mathematical Biology, Mathematical Institute, University of Oxford, Radcliffe Observatory Quarter, Woodstock Road, Oxford, OX2 6GG; Centre for Mathematical Biology, Department of Mathematical Sciences, University of Bath, Claverton Down, Bath, BA2 7AY

## Abstract

In this work we study the effect of domain growth on spatial correlations in agent populations containing multiple species. This is important as heterogenous cell populations are ubiquitous during the embryonic development of many species. We have previously shown that the long term behaviour of an agent population depends on the way in which domain growth is implemented. We extend this work to show that, depending on the way in which domain growth is implemented, different species dominate in multispecies simulations. Continuum approximations of the lattice-based model that ignore spatial correlations cannot capture this behaviour, while those that explicitly account for spatial correlations can. The results presented here show that the precise mechanism of domain growth can determine the long term behaviour of multispecies populations, and in certain circumstances, establish spatially varying species densities.

## 1 Introduction

Heterogeneous cell populations are widespread throughout biology. Obvious examples include the immune system [1], the brain [2], and the heart [3]. Tumours are often composed of cells that are not phenotypically identical, an important factor that reduces the efficacy of many drug treatments [4]. Cranial neural crest stem cells, a subset of a migratory cell population that give rise to a diverse lineage, exhibit ‘leader’ or ‘follower’ phenotypes during their collective cell migration [5–7]. Similarly melanoblasts, another neural crest stem cell subset, and keratinocytes, simultaneously populate the dorsal lateral epithelium during embryonic development [8].

Spatial structure in cell populations is known to be important for their function and development. For instance, in melanoblasts spatial correlations between migrating cells are hypothesised to underpin pigmentation patterns [8]. Spatial structure is often established by cell proliferation, as a new cell is naturally close to its parent cell following division. Important examples of this are tumour development [9] and the growth of the cerebral cortex [2]. Spatial correlations between cells can also be indicative of different types of cell-cell interactions, such as adhesion or repulsion [10–13]. Importantly, many of the aforementioned examples of heterogenous cell populations in which spatial structure is important are associated with growing tissues, either during embryonic development [2, 5–8], or in pathological scenarios [3, 4]. Therefore, it is important to be able to include domain growth in models of cell populations containing multiple species where spatial structure plays a significant role [5–8, 14, 15].

In this work we examine the effects of domain growth on the evolution of spatial correlations between agents (where agents represent cells) in individual-based models (IBMs). To do so we employ an agent-based, discrete random-walk model on a two-dimensional square lattice with volume-exclusion. We have previously shown that the way in which domain growth is implemented in an IBM can alter the behaviour of a population of identical agents [16, 17]. Therefore, we hypothesised that the way in which domain growth is implemented in a simulation with multiple agent species could change the dominant agent species. We also reasoned that a standard mean-field approximation (MFA) would be insufficient to capture the behaviours exhibited by the IBM, and the MFA would require correction by the inclusion of spatial correlations in the form of a system of ordinary differential equations (ODEs) to accurately approximate the IBM results. This has previously been shown in scenarios without growth. For example, Markham et al. [14] demonstrated the necessity of including the effect of spatial correlations in continuum models to accurately predict the dominant species in a multispecies context.

The outline of this work is as follows: we introduce a two-dimensional IBM and two distinct growth mechanisms in Section 2.1. We then define the individual and pairwise density functions, and a derive a system of equations (referred to as a correlation ODE model) describing the evolution of the individual and pairwise density functions for multiple species on a growing domain in Sections 2.2-2.3. In Section 3 we test the accuracy of the correlation ODE model by comparing it with the standard MFA and IBM results for multispecies simulations. We then demonstrate that the precise details of the implementation of domain growth can affect agent population fates; a species that dominates in one growth regime might not dominate in the other. As far as we are aware this is the first time it has been reported that the particular details of the growth regime can change the competition outcome between two species. We also demonstrate that the MFA is unable to accurately capture the effects of domain growth in the IBM, whereas the correlation ODE models that include the effect of spatial correlations do. Finally, we examine some biologically motivated examples of non-uniform domain growth in our IBM in Section 3.2, and show that non-uniform domain growth can cause spatially varying species densities in multispecies agent populations that depend on the growth mechanism implemented. We conclude in Section 4 with a discussion of our results.

## 2 Model

In this section we first introduce the IBM and the two domain growth mechanisms we employ throughout this work. We then derive equations describing the evolution of the individual and pairwise density functions in the IBM for both growth mechanisms. The inclusion of the effect of agent motility, proliferation, and death, in the density functions has been previously presented [14, 18, 19]^1^

### 2.1 IBM and domain growth mechanisms

The IBM is simulated on a two-dimensional square lattice with lattice spacing ∆ = 1 [20] and size *N_x_*(*t*) by *N_y_*(*t*), where *N_x_*(*t*) is the number of lattice sites in a row and *N_y_*(*t*) is the number of sites in a column. Initially, all simulations are performed with periodic boundary conditions.

Each agent is assigned to a lattice site, from which it can move or proliferate into an adjacent site. If an agent attempts to move into a site that is already occupied, the movement event is aborted. Similarly, if an agent attempts to proliferate into a site that is already occupied, the proliferation event is aborted. Processes in which only one agent is allowed per site are often referred to as exclusion processes [20]. Time is evolved continuously, in accordance with the Gillespie algorithm [21], such that movement, proliferation and growth events are modelled as exponentially distributed ‘reaction events’ in a Markov chain. Throughout this work we only present examples with two species, and these species are referred to as *A* and *B*. However, all the results presented in this work are easily extendable to scenarios containing more than two species [14]. Attempted agent movement or proliferation events occur with rates 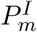 or 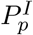 per unit time, respectively, where *I* denotes the species type. For example, 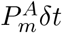 is the probability of an individual agent of species *A* attempting to move in the next infinitesimally small time interval *δt*. Death events occur with rate 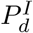 per unit time and result in the removal of an agent from the lattice.

Both growth mechanisms we employ are stochastic [22]: the insertion of new lattice sites occurs with positive rate constants *P_gx_* and *P_gy_* per unit time for growth in the *x* (horizontal) and *y* (vertical) directions, respectively. That is, an individual lattice site undergoes a growth event in the *x* direction with rate *P_gx_*. Our growth mechanisms are designed to represent the growth of an underlying tissue upon which a cell population is situated. Examples of cell populations that are situated on top of growing tissues can be readily found in biological systems. For instance, cell populations such as neural crest stem cells are known to migrate through growing tissues, such as the epidermis, during embryonic development [5, 8, 15]. In growth mechanism 1 (GM1) when a ‘growth event’ occurs along the *x*-axis (horizontal axis in Fig. 1 (a)), one new column of sites is added at a position selected uniformly at random. In growth mechanism 2 (GM2) when a ‘growth event’ occurs along the *x*-axis (see Fig. 1 (b)), for each row, one new site is added in a column that is selected uniformly at random. Importantly, when a growth event occurs, the site selected for division is moved one spacing in the positive horizontal direction along with its contents (i.e. an agent or no agent, an agent is symbolised by a black circle in Fig. 1). The new lattice site is empty, and the contents of all other lattice sites remain unaffected. Growth in the *y* direction is implemented in an analogous manner to the *x* direction for both growth mechanisms. We chose these growth mechanisms as they are significantly different to each other, and both may have biological relevance [23–25]. Furthermore, both of these growth mechanisms can be used to implement any form of isotropic growth in our IBM, and are adaptable to three spatial dimensions [22]. Finally, it is important to note that both growth mechanisms give rise to the same overall growth rate when implemented with the same rate constants.

**Figure 1:**
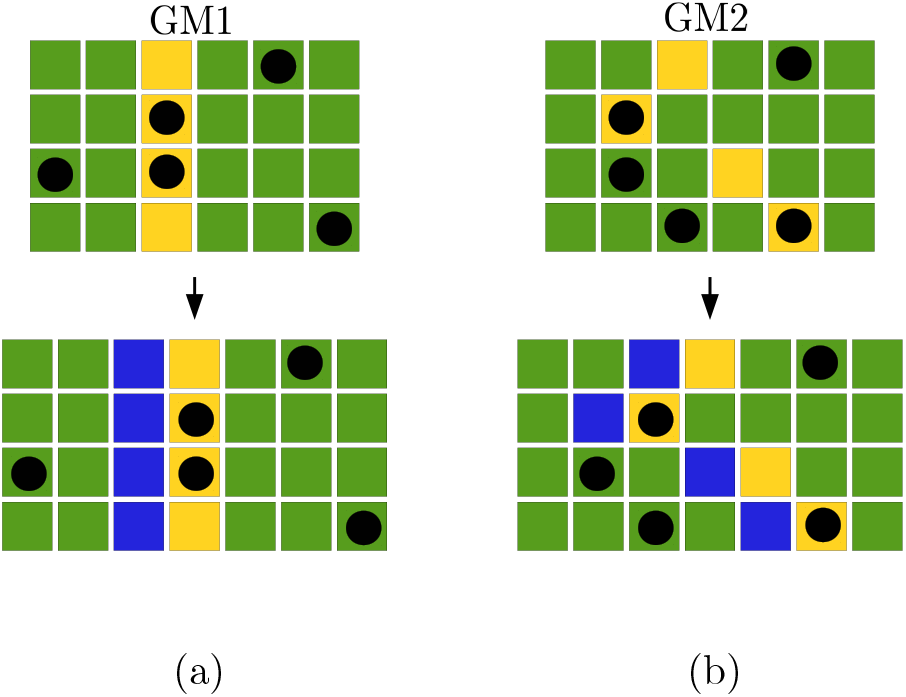
(Colour online). Before and after the growth events for both (a) GM1 and (b) GM2, in which growth is along the *x*-axis for a two-dimensional lattice. In each row the yellow (light grey) site has been chosen to undergo a growth event. Following this the yellow (light grey) site is moved to the right with its contents, for instance an agent (represented by a black cell). The blue (dark grey) sites are the new lattice sites and are always initially empty. The contents of all the other sites remain unaffected, although in some cases their neighbouring sites will change.

Throughout this work we employ homogenous initial conditions in our IBM (when our initial condition is averaged over many repeats). That is, our initial distribution for both species is achieved by populating a certain number of sites uniformly at random. An occupied site is indicated by *A* or *B*, and an unoccupied site is indicated by 0. This means the normalised average agent density for species *A* on the two-dimensional domain is

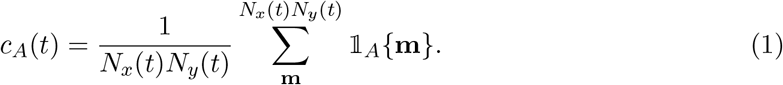
Here 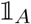 is the indicator function for species *A* (i.e. 1 if species *A* occupies site **m**, and 0 if it does not). An analogous equation holds for species *B*.

### 2.2 Individual density functions

We now derive the evolution equations for the individual density functions. To begin with, we only include the effects of domain growth on the density functions (the details of how to include the effects of agent motility, proliferation, and death, in the density functions are demonstrated in the supplementary information Section S1).

We define the individual density functions, 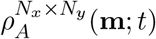, as the probability that site **m** is occupied by an agent *A* at time *t* on a domain of size *N_x_*(*t*) × *N_y_*(*t*), where **m** is the vector (*i, j*), with *i* indexing the row number of a lattice site, and *j* indexing the column number of a lattice site^2^. For instance, (2, 3) would be the lattice site situated in the second row and the third column of the lattice. Similarly, 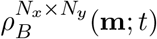 is defined as the probability that site **m** is occupied by an agent *B* at time *t* on a domain of size *N_x_*(*t*) *× N_y_*(*t*).

The following derivation for the evolution of individual density functions is the same for GM1 and GM2 (and for species *A* and *B*)^3^. Therefore we only derive the equation for the evolution of the individual density functions in the case of species *A*. The sum of the individual density functions on a domain of size *N_x_*(*t* + *δt*) × *N_y_*(*t* + *δt*) at [*t* + *δt*) for species *A* can be written in terms of the individual density functions at time *t*:

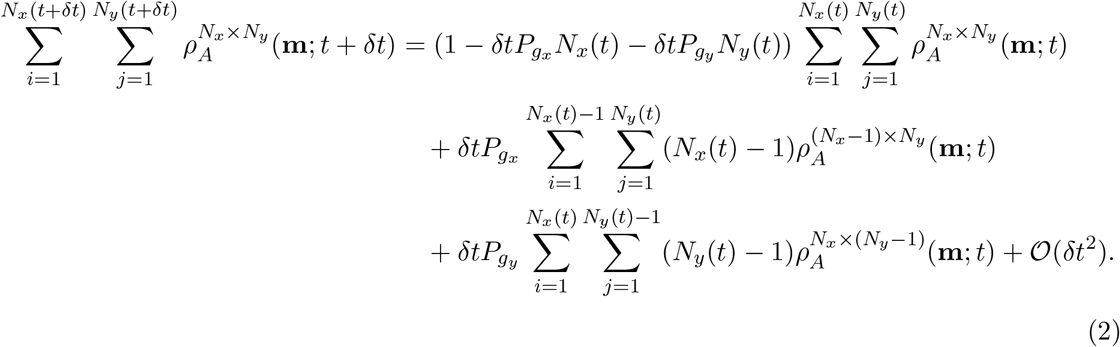

The terms of the right-hand-side (RHS) of Eq. (2) correspond to the following events: i) no growth event occurs in [*t, t* + *δt*); ii) a growth event occurs in the horizontal (*x*) direction in [*t, t* + *δt*); and iii) a growth event occurs in the vertical (*y*) direction in [*t, t* + *δt*). As the initial conditions are, on average, spatially uniform we can assume translational invariance for the probability of an agent occupying a site throughout. By this we mean

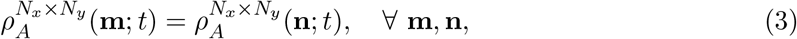

where **n** indexes any other site on the domain^4^.

Equation (3) allows us to rewrite Eq. (2) as

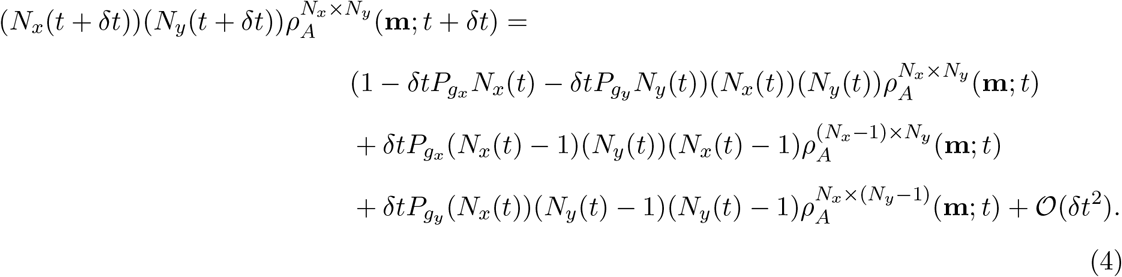

Equation (4) can then be simplified to obtain

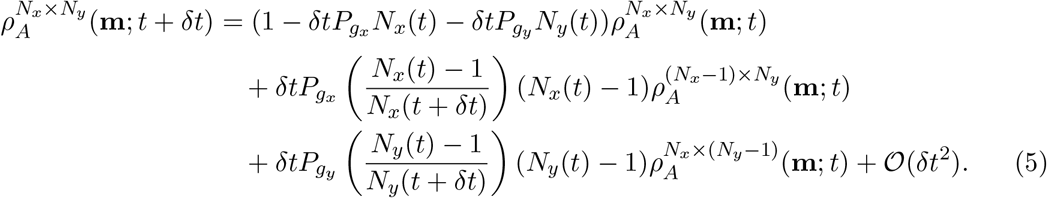

Rearranging Eq. (5) and taking the limit as *δt →* 0 we arrive at the ODE

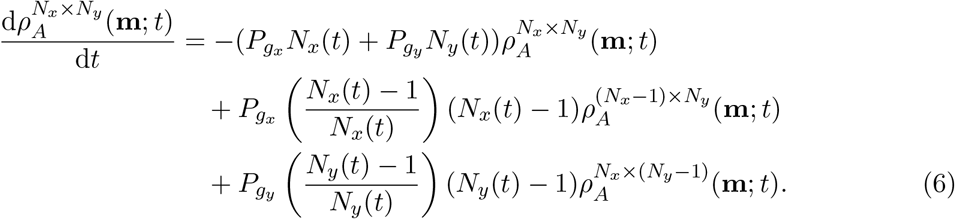

If we make the approximation 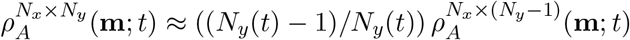 and 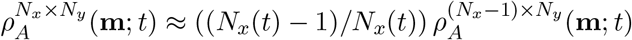 in Eq. (6) we obtain

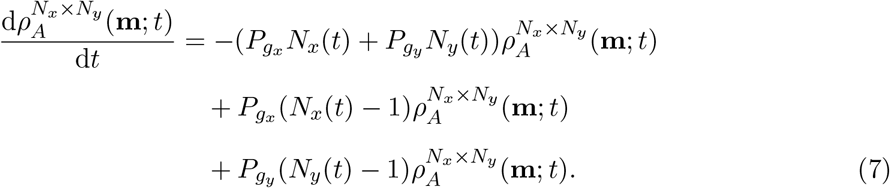

This approximation has been previously published [26], and reasonably implies that domain growth ‘dilutes’ the agent density [16] (we present analysis of the accuracy of this approximation in Section S2 of the supplementary information). Finally, we simplify Eq. (7) to obtain

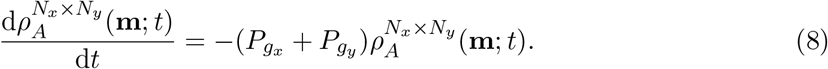

Equation (8) is a single equation that describes how exponential domain growth affects the evolution of the individual density functions for species *A*. It is important to note that Eq. (8) describes how *exponential* domain growth affects the evolution of individual density functions because we have defined *P_gx_* and *P_gy_* as constants. It is straightforward to derive equations for linear and logistic domain growth analogous to Eq. (8) if required.

In the course of the following derivation it will be useful to write the pairwise density functions in terms of the distances between sites, therefore we shall rewrite the individual density functions as 
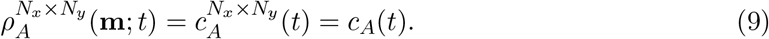

If we substitute Eq. (9) into Eq. (8) we obtain 
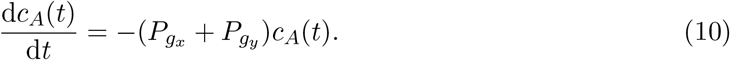
A comparison between Eqs. (6) and (10) demonstrates that the approximation we have employed reduces an infinite system of equations describing the evolution of the macroscopic agent density on a growing domain, into a single first-order linear ODE that is trivially solvable. To include the effects of agent proliferation and motility in Eq. (10) we first need to define the pairwise density functions.

### 2.3 Pairwise density functions

Figure 2 displays two configurations of two agents, which we will term (a) colinear and (b) diagonal. The distance between sites is measured from their centres, as illustrated in Fig. 2 (b).

**Figure 2:**
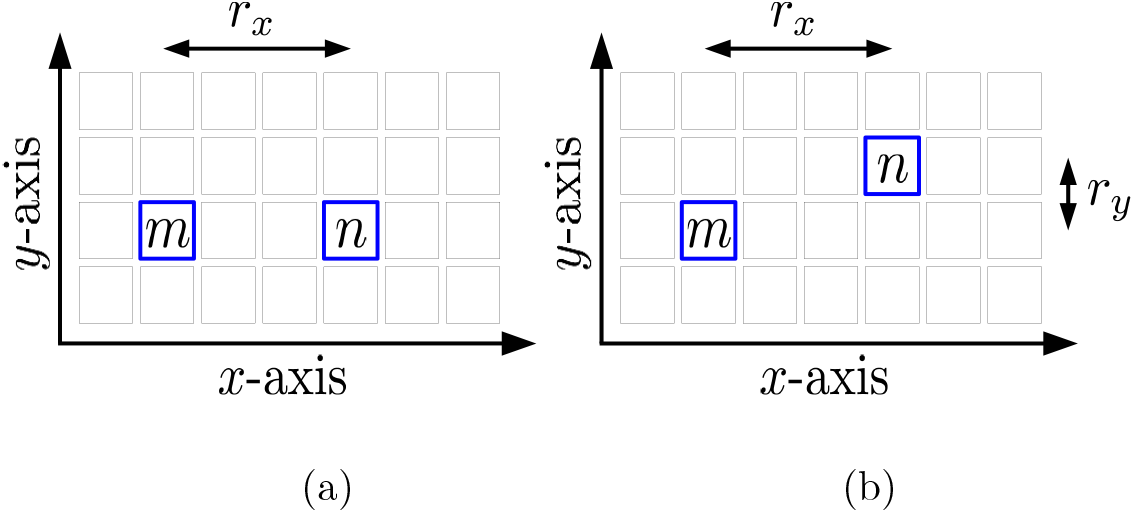
The two types of configuration of lattice sites: (a) colinear and (b) diagonal. The lattice sites in question are labelled **m** and **n** and bordered by blue (thicker line). In (a), two colinear lattice sites share the same row but not the same column (or vice versa). In (b), two lattice sites are diagonal, meaning they do not share the same row or column. *r_x_* is the distance between two lattice sites in the horizontal direction, *r_y_* is the distance between two lattice sites in the vertical direction. In (b) *r_x_* = 3 and *r_y_* = 1.

As can be seen in Fig. 2, *r_x_* is the distance between two lattice sites in the horizontal direction, and *r_y_* is the distance between two lattice sites in the vertical direction.

We define the *auto*-correlation pairwise density functions, 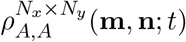, as the probability sites **m** and **n** are both occupied by species *A* at time *t* on a domain of size *N_x_*(*t*) *× N_y_*(*t*) (where **m** ≠ **n**). Similarly, the auto-correlation pairwise density function 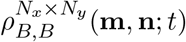 is defined as the probability sites **m** and **n** are both occupied by species *B* at time *t* on a domain of size *N_x_*(*t*) *× N_y_*(*t*). The *cross*-correlation pairwise density function, 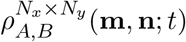, is the probability sites **m** and **n** are occupied by species *A* and *B*, respectively, at time *t* on a domain of size *N_x_*(*t*) *× N_y_*(*t*). We now rewrite the pairwise density functions in terms of the displacement vector between lattice sites, that is 
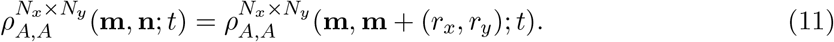

As the initial conditions in the IBM are, on average, spatially uniform we are able to assume translational invariance for the probability of two sites a given distance apart being occupied. This means that the pairwise density function can be written as a function of the displacement between two lattice sites, (*r_x_, r_y_*). Therefore, we will further simplify our notation and write 
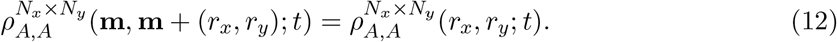

#### 2.3.1 Agent motility, proliferation and death

The inclusion of agent motility, proliferation and death in Eq. (10) has been outlined previously [14, 18, 19]. Therefore, we refer the reader to the supplementary information (Section S1) for details of how to include the effects of agent motility, proliferation, and death in Eq. (10), and simply state the result for the individual density functions in the main text. The evolution of the individual density function for a motile and proliferating species *A* on a growing domain is

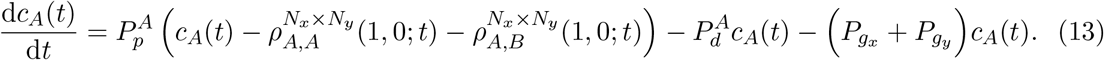

An analogous equation exists for species *B*. As can be seen from Eq. (40), the inclusion of agent proliferation means that pairwise density functions are now present in the equations for the evolution of the individual density functions, which is not the case without agent proliferation (Eq. (10)). It is important to stress that we can combine proliferation, death, and growth terms as we do in Eq. (40), as these terms are independent of each other in the derivation of Eq. (40) (see the supplementary information Section S1 for further details).

#### 2.3.2 Growth mechanism 1

We now derive the equations for the evolution of the pairwise density functions for GM1 domain growth. We do not include agent migration, proliferation, or death in the following derivation for the purposes of clarity. The inclusion of the effects of agent migration, proliferation, and death in the equations for the evolution of the pairwise density functions has been described before [14, 18, 19] (the details of how to do so can be found in the supplementary information Section S1).

##### Colinear component

We begin with the colinear component of the equations for the evolution of the pairwise density functions, that is, the scenario in which the lattice sites in question share the same column or row, as depicted in Fig. 2 (a). The following derivation is the same for both auto-correlation pairwise density functions, 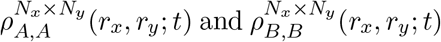, and the cross-correlation pair-wise density function, 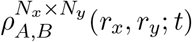. Therefore, we only derive the evolution of the pairwise density functions for species *A*. For agents colinear in the horizontal direction, that is, *r_y_* = 0, the evolution of the auto-correlation pairwise density functions for species *A* with GM1 is

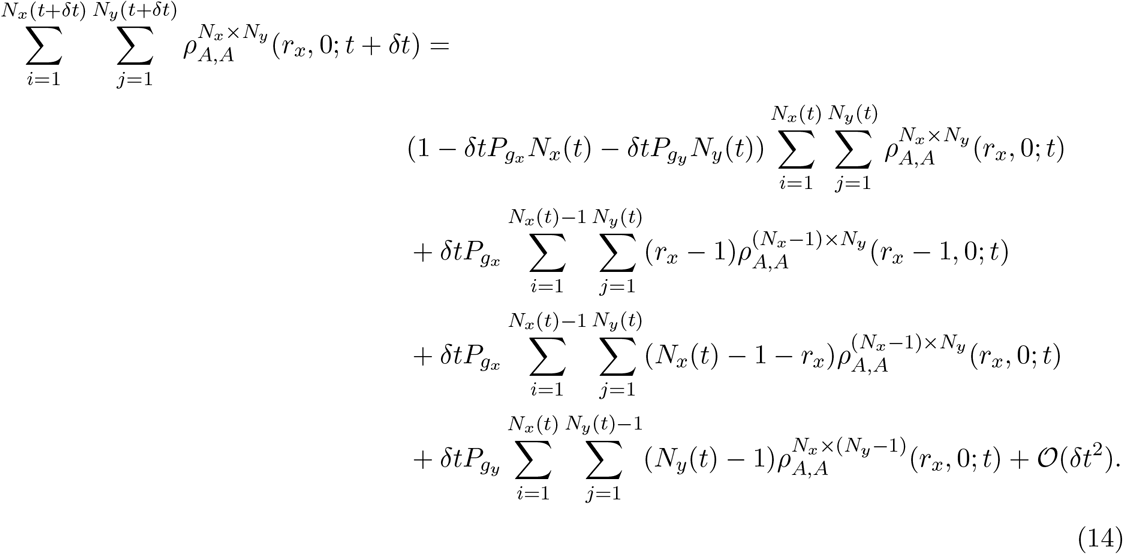

The terms on the RHS represent the probabilities that: i) no growth event occurs in [*t, t* + *δt*); ii) a growth event occurs in the horizontal direction between agents (*r_x_ −* 1, 0) apart, moving them (*r_x_,* 0) apart on a domain of size *N_x_*(*t* + *δt*) *× N_y_*(*t* + *δt*) at [*t* + *δt*); iii) a growth event occurs in the horizontal direction at a site that is not in between agents (*r_x_,* 0) apart, meaning that they remain (*r_x_,* 0) apart but now on a domain of size *N_x_*(*t* + *δt*) *× N_y_*(*t* + *δt*) at time [*t* + *δt*); and iv) a growth event occurs in the vertical direction (as the sites are horizontally colinear in this a GM1 growth event cannot change the displacement between them).

Similarly, the evolution of the auto-correlation pairwise density functions for agents colinear in the vertical direction (that is, *r_x_* = 0) is

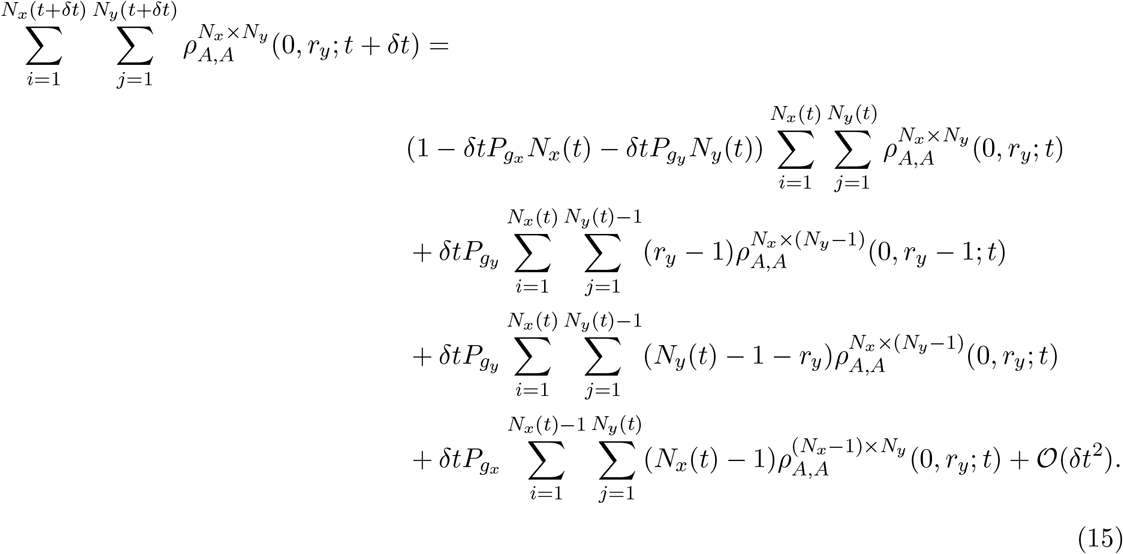

We can simplify Eq. (14) to obtain

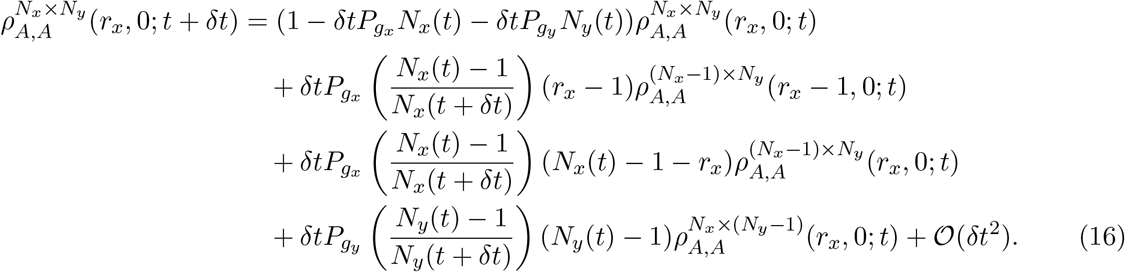

If we apply the approximations^5^ 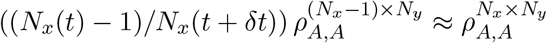 and 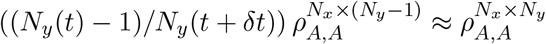 to Eq. (16) we obtain

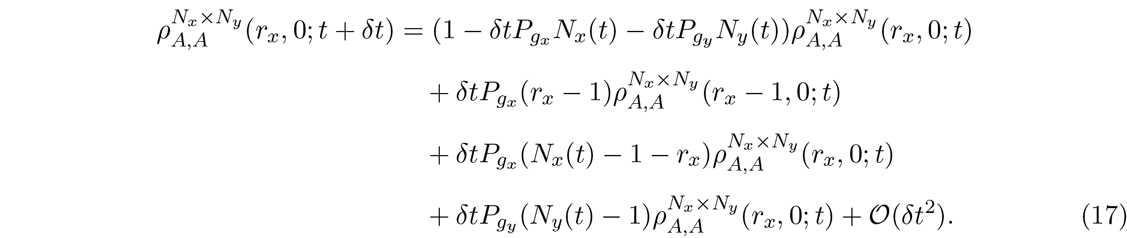

Rearranging Eq. (17) and taking the limit as *δt →* 0 we arrive at

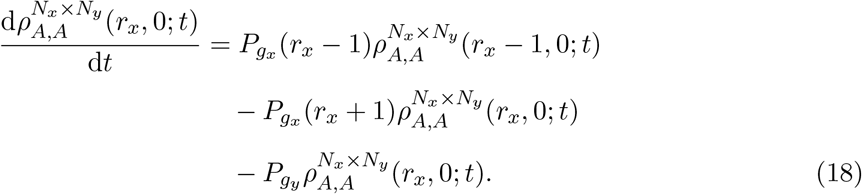

The equivalent equation for sites colinear in the vertical direction (see Eq. (15)) is

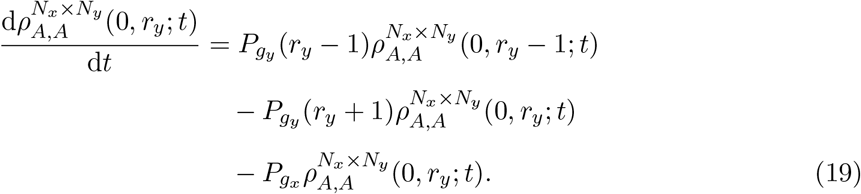

##### Diagonal component

For the diagonal component, that is, *r_x_, r_y_* ≠ 0, we have, by similar reasoning,

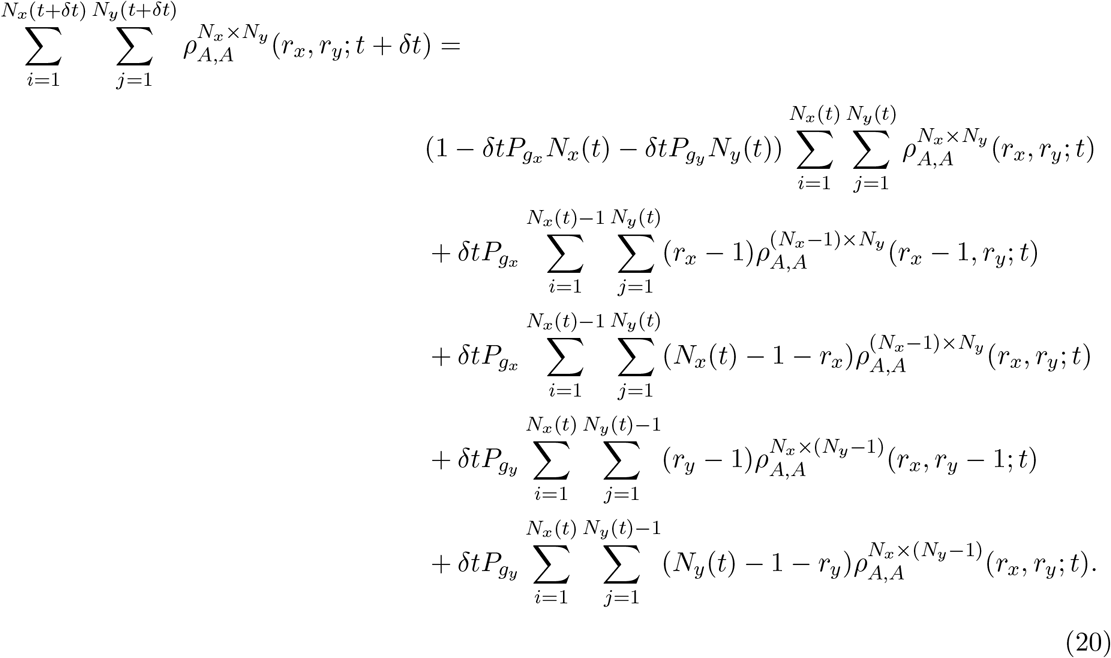

If we follow the same procedure as for the colinear component we obtain

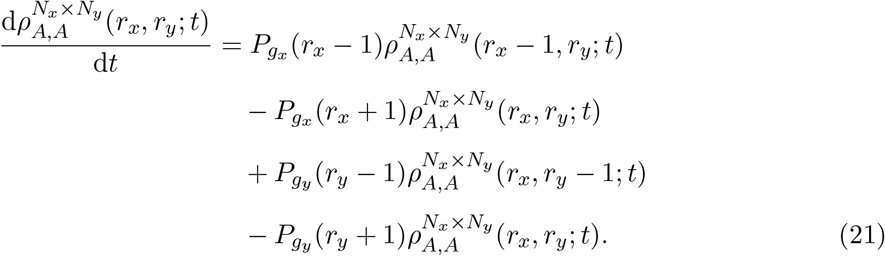

#### 2.3.3 Growth mechanism 2

For the derivation of the pairwise density functions for GM2 we refer the reader to the supplementary information (Section S3) and simply state the results in the main text. The evolution equation for the colinear component (horizontally colinear) for GM2 is

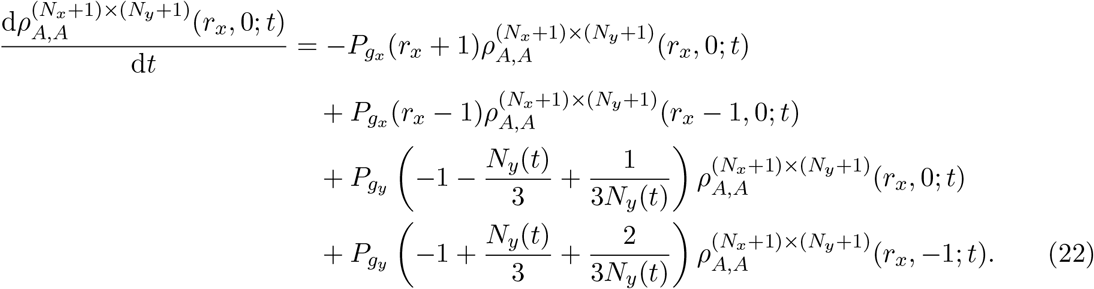

An analogous equation exists for the vertically colinear component for GM2. The diagonal component for GM2 is

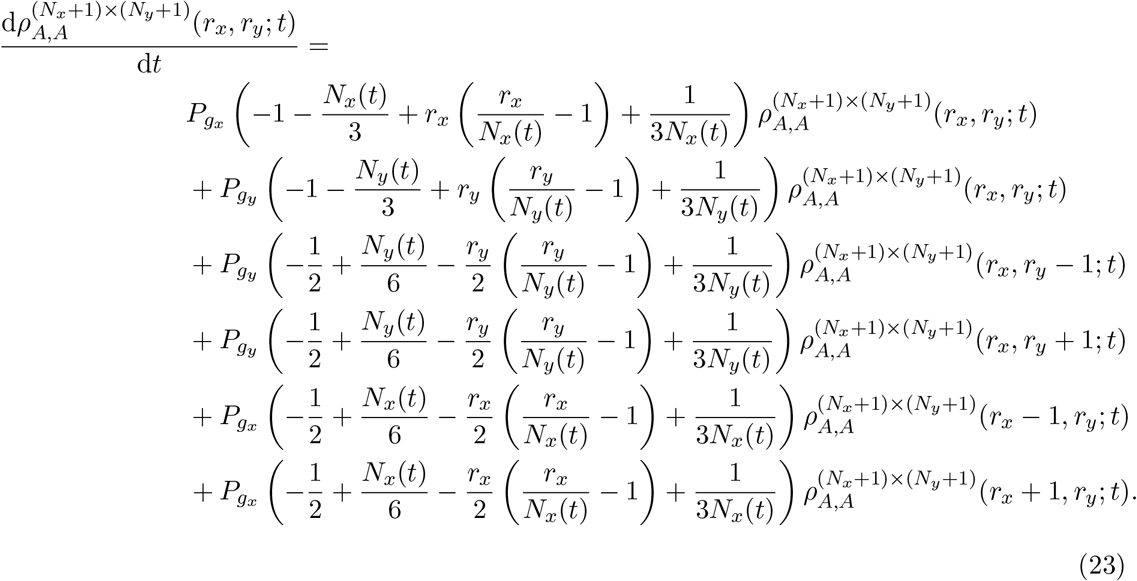

If we compare Eqs. (18) and (21)-(23) it is apparent that the length of the domain influences the evolution of the pairwise density functions in the case of GM2, but not in GM1.

## 3 Results

We present our results in terms of correlation functions [27–31] in order to simplify the visualisation of results, and allow the results presented here to be easily related to other research in this field [14, 18, 19, 32, 33]. The correlation function is defined as

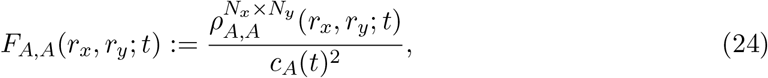
and is simply a measure of the degree to which the occupancies of two lattice sites are independent of one another. Analogous correlation functions exist for auto-correlations in species *B* and for the cross-correlations between species *A* and *B*:

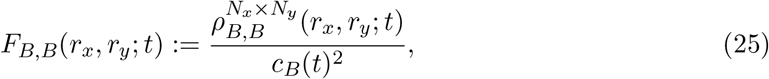
and 
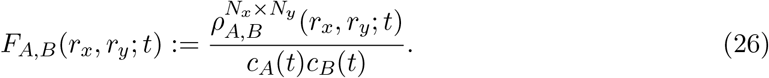

If we substitute Eqs. (24) and (26) into Eq. (40) we obtain 
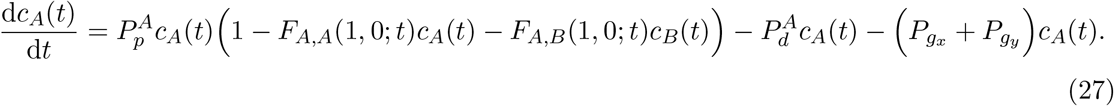

We refer to Eq. (27) as the correlation ODE model. The standard MFA assumes *F_A,A_*(1, 0; *t*) = *F_A,B_* (1, 0; *t*) = 1, that is, spatial correlations between agents are insignificant, and so Eq. (27) becomes
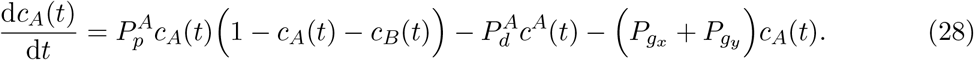

Equation (28) is relevant as it represents the standard MFA often used to model the evolution of the macroscopic density of a cell population [13, 34, 35]. However, in certain scenarios the standard MFA has been shown to be inadequate [18], especially when the spatial structure of a cell population is known to be important. As such, we will compare Eqs. (27)-(28) in the results section.

For our discrete simulations we use a regular square lattice of initial size 100 by 100 lattice sites. The boundary conditions are periodic, and we have an initial uniform random seeding of density 0.05 for each species *A* and *B* (so the total agent density is 0.1). By an initial uniform random seeding it is meant that, on average, the initial conditions of the IBM are spatially uniform for both species. All results presented from the IBM are ensemble averages taken from 500 repeats. To solve Eqs. (10), (18) and (21)-(40) we use MATLAB’s ode15s, with an absolute error tolerance of 10^−12^.

Our initial condition for all simulations entails that all pairwise distances are initially uncorrelated, that is,

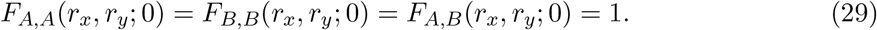

Initially we study the effect of exponential domain growth for both GM1 and GM2 on agent density and spatial correlations in the IBM and correlation ODE model. As previously stated, *N_x_*(*t*) and *N_y_*(*t*) are integers that describe the number of lattice sites in the horizontal and vertical directions, respectively. However, as results from the IBM are ensemble averages we replace *N_x_*(*t*) and *N_y_*(*t*) with their continuum analogues *L_x_*(*t*) and *L_y_* (*t*), respectively. This substitution of *N_x_*(*t*) and *N_y_*(*t*) with *L_x_*(*t*) and *L_y_* (*t*) avoids jump discontinuities in the numerical solutions of Eqs. (21)-(23), which are not present in the averaged IBM results. For exponential domain growth *L_x_*(*t*) evolves according to

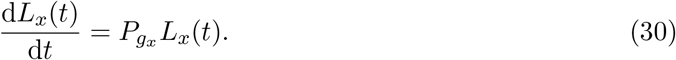

An analogous equation exists for *L_y_*(*t*).

We also study the effect of linear and logistic domain growth for both GM1 and GM2 on agent density and spatial correlations in the IBM and correlation ODE model. For simulations with logistic domain growth the individual density function for species *A* evolves according to

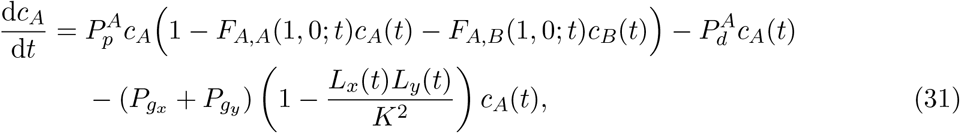
where *K*^2^ is the carrying capacity and *L_x_*(*t*) evolves according to

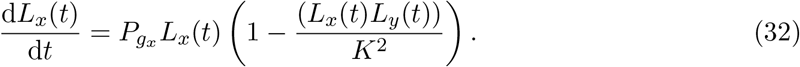

An analogous equation exists for *L_y_*(*t*), and for all simulations presented here *K* = 300. Finally, the individual density function for species *A* with linear domain growth evolves according to

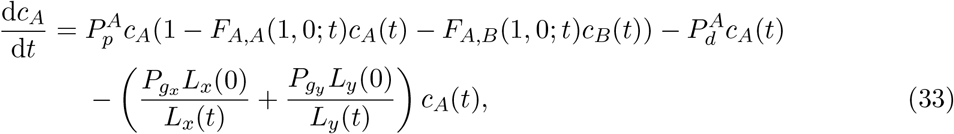
where *L_x_*(*t*) evolves according to 
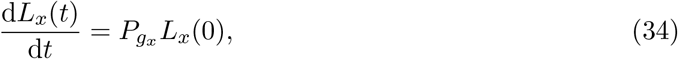
that is, linear growth. We also rescale time to allow for ease of comparison between simulations with different parameters: 
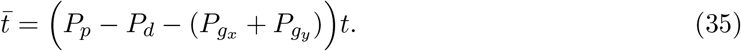

### 3.1 Uniform domain growth

In Fig. 3 we recapitulate results from [14]^6^. We see that in the case of an IBM with a non-growing domain and two species a more motile, slower proliferating species (species *A*) can dominate over a less motile, faster proliferating species (species *B*) given a specific parameter regime. This is the case on a non-growing domain without agent death as can be seen in Fig. 3 (a), and is augmented with agent death as evident in Fig. 3 (b), whereby species *B* eventually goes extinct. Importantly, the standard MFA (Eq. (28)) is not able to accurately approximate the IBM results in either case, whereas the correlations ODE model is able to.

**Figure 3:**
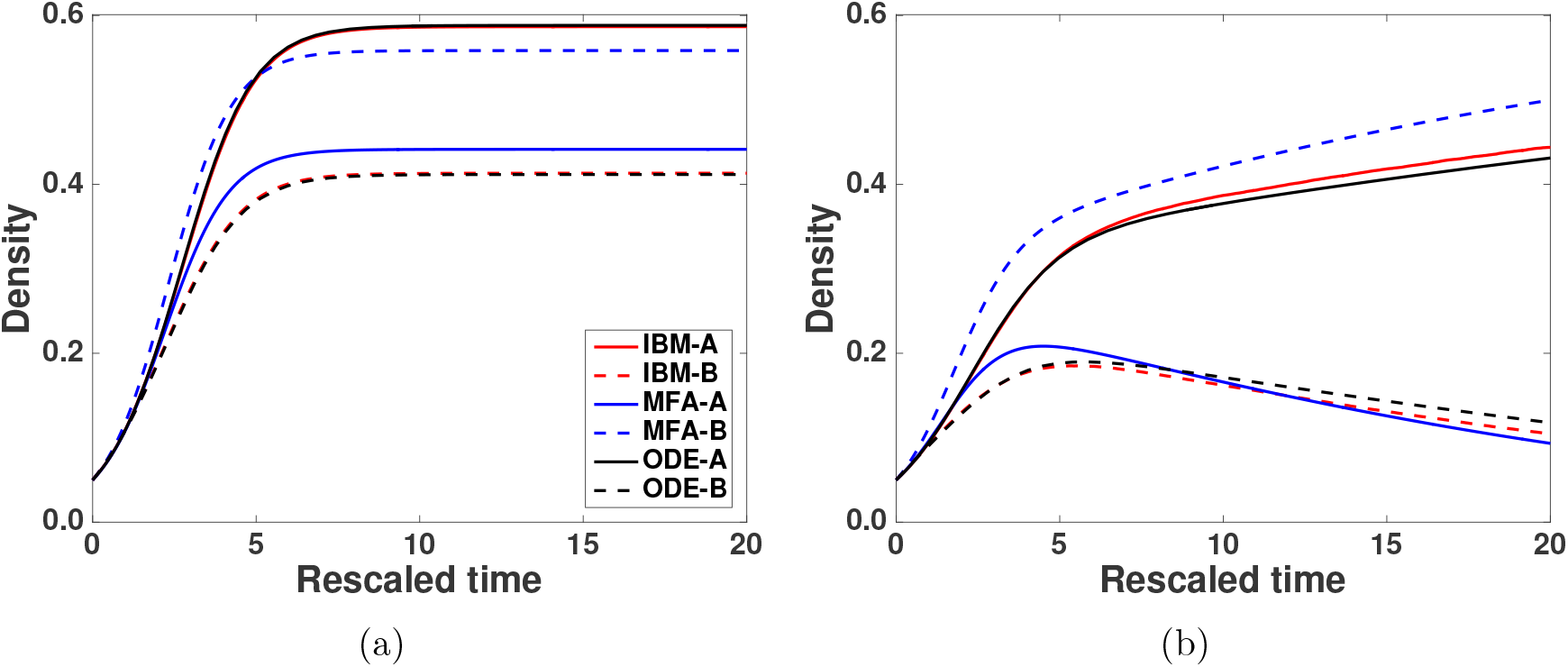
(Colour online): Including the effects of pairwise correlations renders the correlations ODE model (Eq. (27)) able to accurately approximate the averaged results from the IBM, whereas the standard MFA (Eq. (28)) cannot. In (a) the parameters are 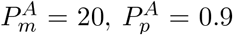, 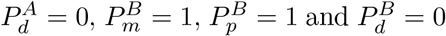. In (b) the parameters are 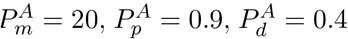, 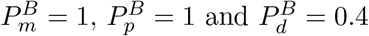.

Figure 4 (a) shows that domain growth implemented via GM1 has a similar effect on agent density as agent death in Fig. 3 (b), allowing species *A* to dominate (although species *B* does not become extinct in this case as there is no agent death). In Fig. 4 (b) we can see that domain growth implemented via GM2 has the opposite effect, and enables species *B* to dominate. This is because GM2 breaks up colinear correlations (correlations between agents that share the same row or column) at a rate proportional to the size of the domain. This means species *B*, which is more affected by colinear correlations due to its higher proliferation and lower motility rates compared to species *A*, increases in ‘fitness’ as the domain grows. Importantly, the standard MFA is not able to capture the GM1 results, whereas the correlations ODE model is. In the GM2 scenario, the MFA ultimately predicts the correct dominant species, but the correlations ODE model more accurately predicts the temporal evolution of the system to 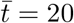.

**Figure 4:**
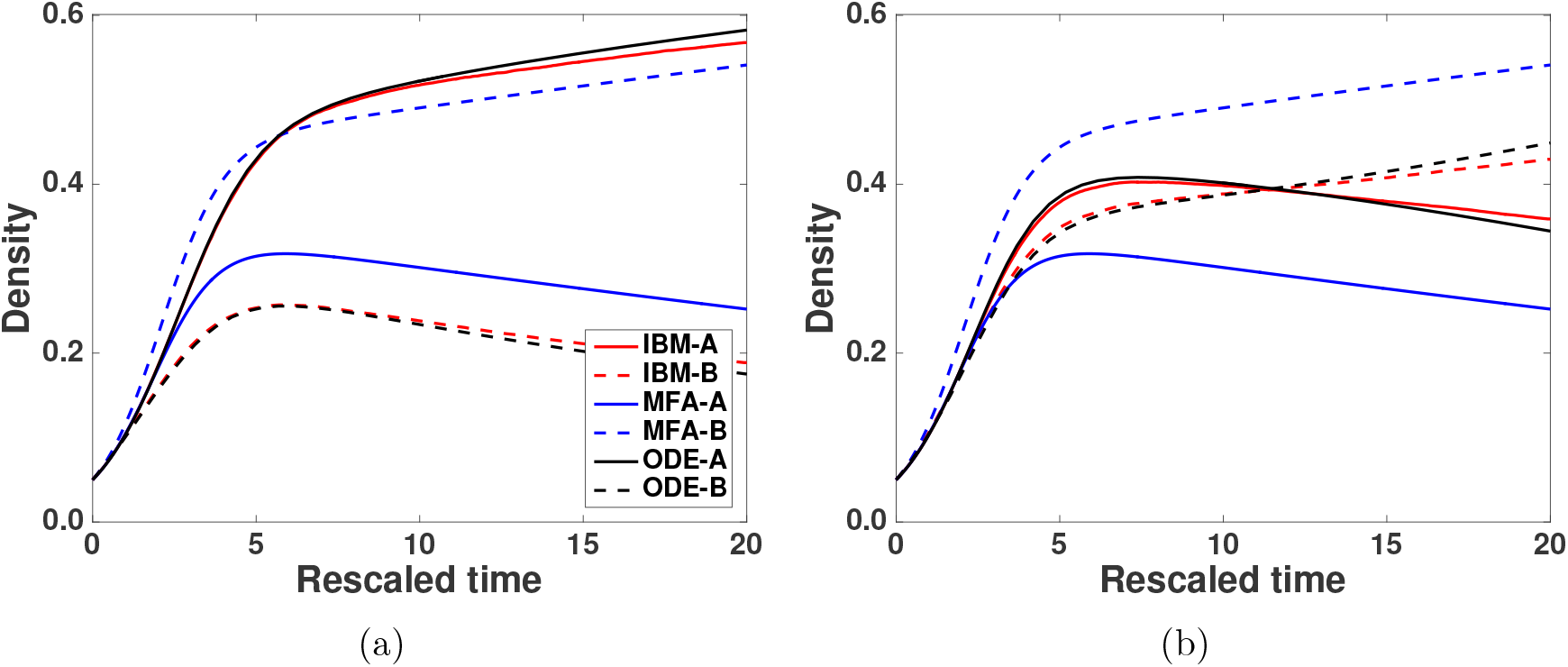
(Colour online): Including the effects of pairwise correlations renders the correlations ODE model (Eq. (27)) able to accurately approximate the averaged results from the IBM, whereas the standard MFA (Eq. (28)) cannot. The parameters for both panels (a) GM1 and (b) GM2 are 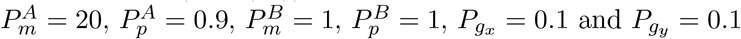.

In Fig. 5 it can be seen that the inclusion of domain growth has a different effect on spatial correlations depending on the type of growth mechanism implemented. We see in Fig. 5 a good agreement between the spatial correlations computed from the correlation ODE model and those calculated from the IBM. For GM1 in Fig. 5 (a) the species *A* auto-correlations decrease as species *A* begins to dominate (long term behaviour). This is because as species *A* begins to dominate its density becomes increasingly spatially uniform. In Fig. 5 (b) we see that species *B* auto-correlations increase as species *B* becomes less spatially uniform, due to proliferation. Meanwhile in Fig. 5 (c) we see that the cross-correlations decrease as species *A* begins to dominate.

**Figure 5:**
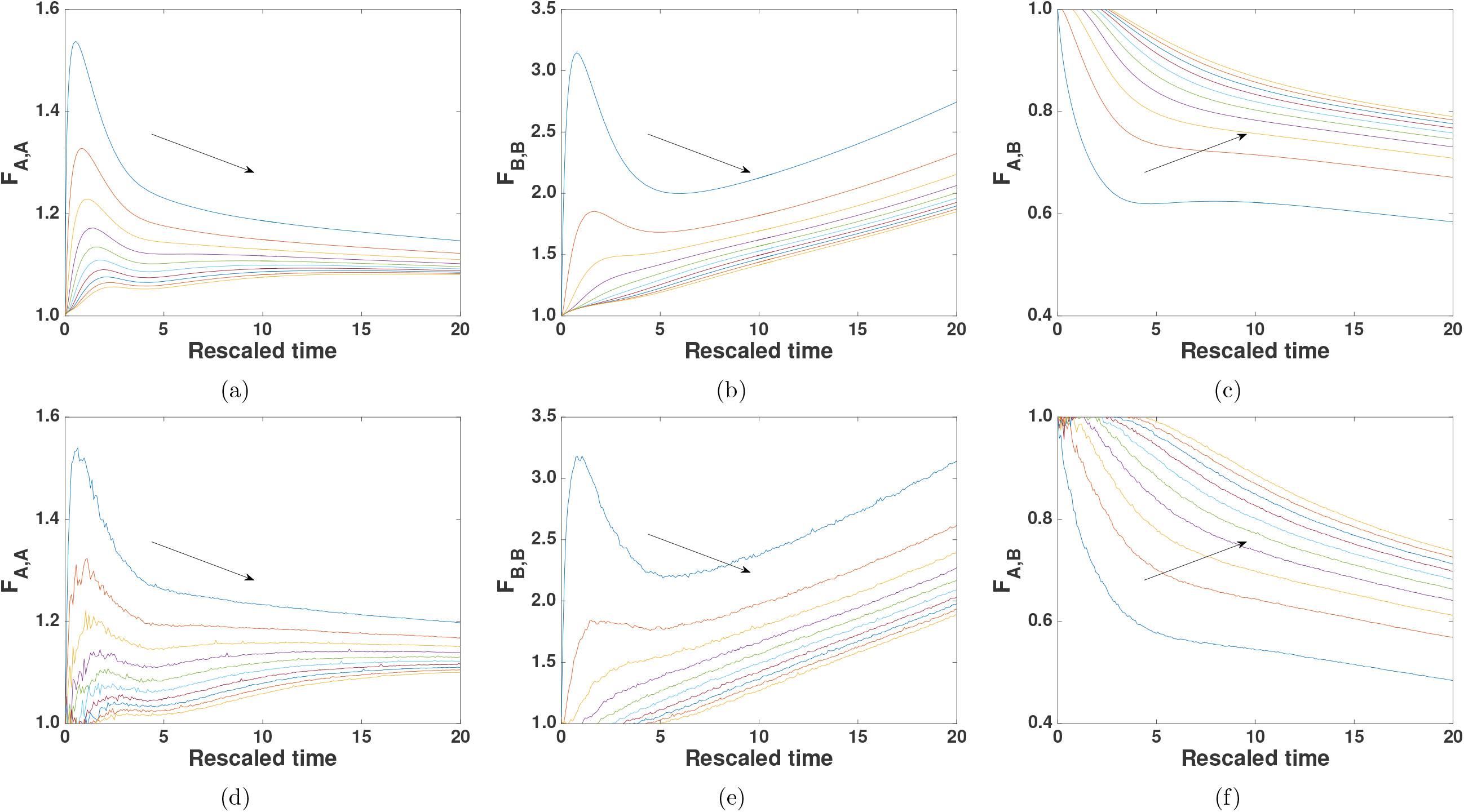
(Colour online): (a)-(c) the evolution of the correlation functions for *F_A,A_*, *F_B,B_* and *F_A,B_* with exponential growth via GM1. The distance plotted increases from ∆ to 10∆ in steps of ∆. Increasing distance is from top to bottom in (a) and (b), i.e. blue line = ∆, red line = 2∆, and so on. In (c) distance increases from bottom to top. In (d)-(f) the correlation functions for *F_A,A_*, *F_B,B_* and *F_A,B_* are calculated from ensemble averages from the IBM. In (a)-(b) and (d)-(e) the auto-correlation functions are greater than or equal to unity, indicating that lattice sites are more likely to be occupied by a particular species if there are others nearby. The lower movement rate of species *B* leads to it having larger auto-correlations, and these auto-correlations grow as species *A* begins to dominate. In contrast, the auto-correlations of species *A* decrease as it begins to dominate. The parameters for all panels are 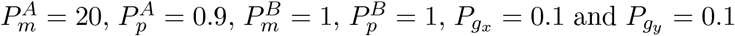.

With GM2 we see that domain growth has a different effect on spatial correlations, as shown in Fig. 6 (d)-(e). Again we see a good agreement between the spatial correlations predicted by the correlation ODE model and those calculated from the IBM. It can be seen that with GM2 the auto-correlations for both species *A* and *B* decrease as the domain grows. In Fig. 6 (f) we see that the cross-correlations between decrease with increasing distance, and note that this is because GM2 breaks up spatial correlations between agents more effectively than GM1.

**Figure 6:**
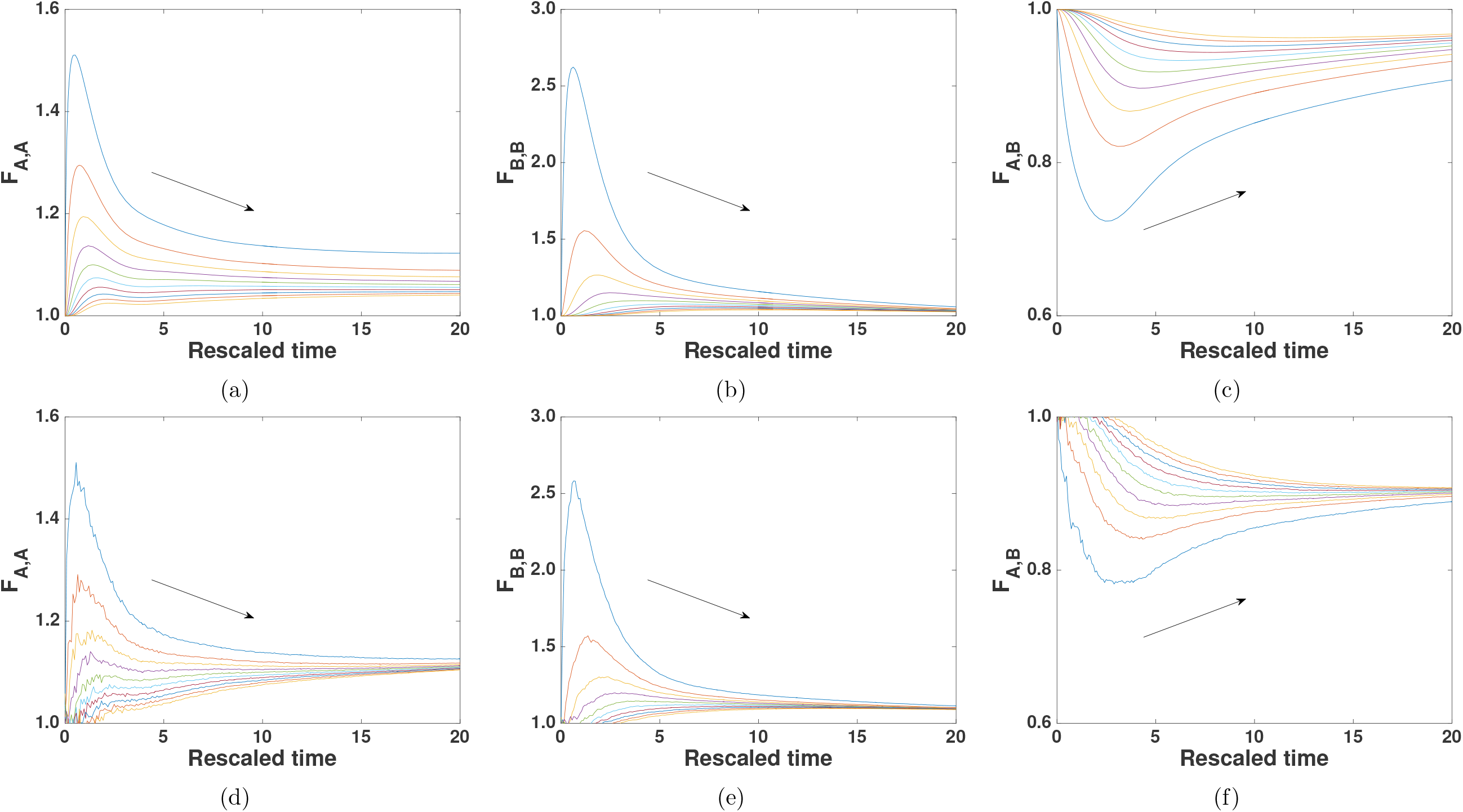
(Colour online): (a)-(c) the evolution of the correlation functions for *F_A,A_*, *F_B,B_* and *F_A,B_* with exponential growth via GM2. In (d)-(f) the correlation functions for *F_A,A_*, *F_B,B_* and *F_A,B_* are calculated from ensemble averages from the IBM. See Fig. 5 for further details. The lower movement rate of species *B* leads to species *B* having larger auto-correlations, and these auto-correlations decrease as species *B* begins to dominate. The parameters for all panels are 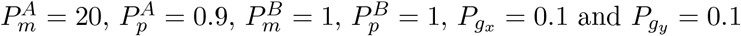.

In Fig. 7 we see that the initial size of the domain influences the evolution of the agent density in the case of GM2, but not in the case of GM1 with exponential domain growth. In the case of GM2, as the initial domain size is increased the evolution of the macroscopic agent densities is accelerated, i.e. species *B* begins to dominate at an earlier time. This is because colinear spatial correlations established by agent proliferation, which affect species *B* more significantly than species *A*, are broken down at a rate proportional to the domain size in GM2. This means the ‘fitness’ of species *B* increases as the domain grows.

**Figure 7:**
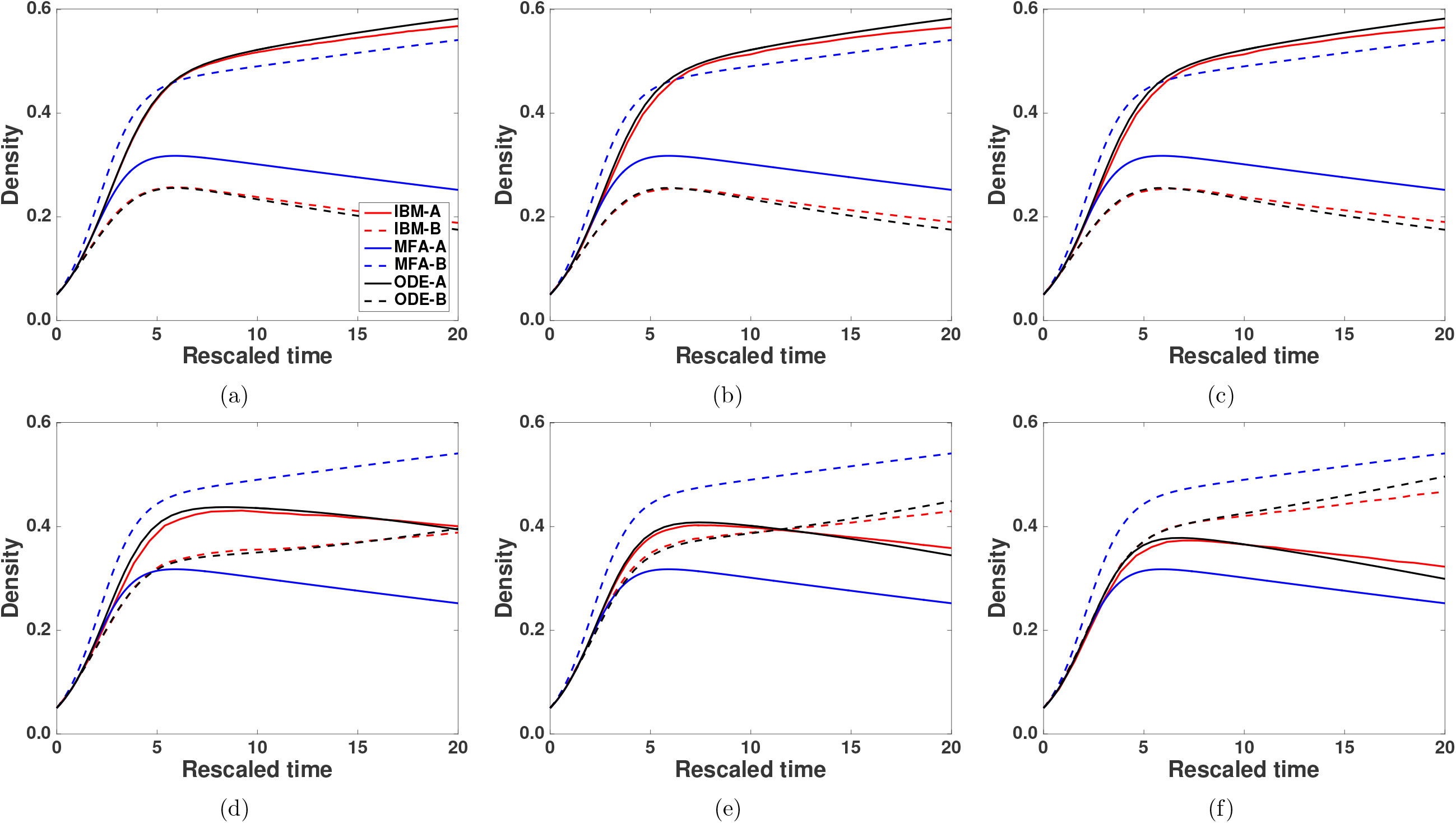
(Colour online): Altering the initial domain size causes the evolution of both species to change in GM2 but not in GM1. As the initial size of the domain is increased the dominance of species *B* in GM2 occurs earlier. GM1: (a) 50 by 50, (b) 100 by 100, (c) 200 by 200. GM2: (d) 50 by 50, (e) 100 by 100, (f) 200 by 200. The parameters for all panels are 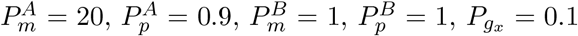

Figure 8 (a)-(b) shows the results from the same two-species scenario with linear domain growth. As before we see that with GM1 species *A* dominates. The correlation ODE model is able to capture this behaviour while the standard MFA is not. In the case of GM2 we see that species *A* initially dominates, but as the domain grows species *B* begins to increase relative to *A* (and the density of species *B* will exceed the density of species *A* at a later time). Finally, Fig. 8 (c) and (d) shows the results from the same two-species scenario with logistic domain growth. In this case we see that species *A* dominates with both GM1 and GM2. This is because the domain stops growing when the domain size carrying capacity is reached.

**Figure 8:**
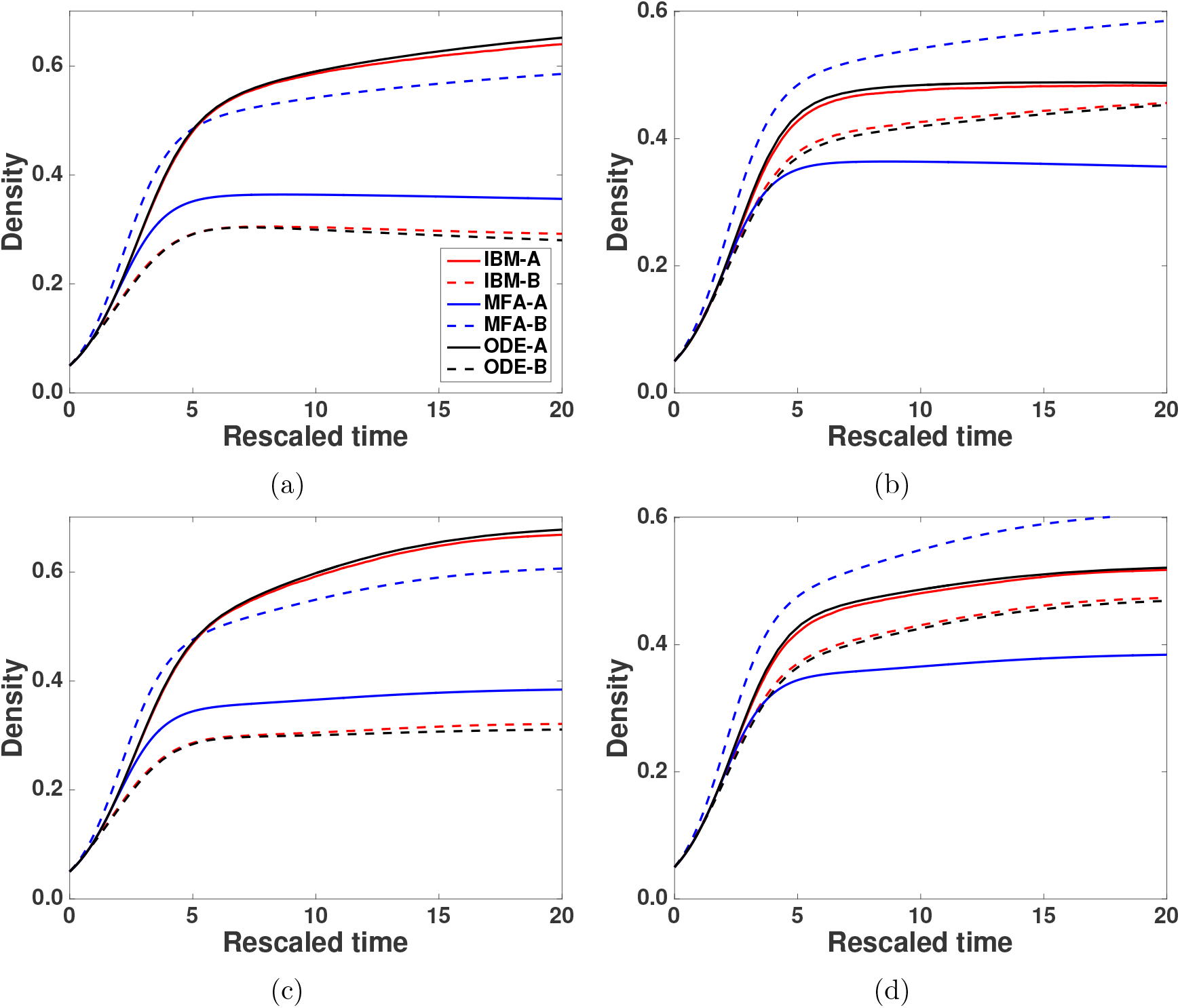
(Colour online): Including the effects of pairwise correlations renders the correlations ODE model (Eqs. (31) and (33)) able to accurately approximate the averaged results from the IBM, whereas the standard MFA cannot. (a) GM1 linear domain growth, (b) GM2 linear domain growth, (c) GM1 logistic domain growth, (d) GM2 logistic domain growth. The parameters for all panels are 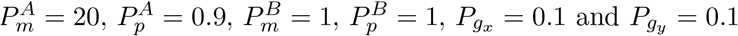.

### 3.2 Non-uniform domain growth

To conclude our results we study some biologically motivated examples of non-uniform domain growth. In these examples we only present results from the IBM. However, we do so aware of the differing effects that GM1 and GM2 have on the evolution of agent density. We hypothesised that, given the same motility and proliferation parameters for species *A* and *B* as in Figs. 3-8, in a two-species scenario non-uniform domain growth could enable species *B* to dominate in a faster growing region of the domain, while species *A* could dominate in a slower growing (or non-growing) region of the domain. This means non-uniform domain growth could lead to spatially variable species densities in simulations containing two species.

We choose two ‘canonical’ examples of domain growth that serve to represent idealised versions of known growth mechanisms in biological systems. The first example we term ‘enteric’. Enteric growth, that is, intestinal growth, is associated with different regions of the intestine growing at different rates [22]. For our enteric example domain growth is again uniform is the vertical direction, while in the horizontal direction ninety percent of the growth events are restricted to the middle third of the *x*-axis (this ‘third’ of the domain is updated throughout the simulation as the IBM domain grows). The other ten percent of growth events are distributed uniformly amongst the two remaining regions.

The second example we term ‘apical’. We use apical to mean domain growth localised to one end of the domain. This type of growth has been observed in root growth and embryonic limb development [36, 37]. For our apical example domain growth is uniform in the vertical direction, while in the horizontal direction growth is restricted to the second half of the *x*-axis (this ‘half’ of the domain is updated throughout the simulation as the IBM domain grows).

For both of these growth mechanisms we implement no-flux boundary conditions in the *x* direction, and periodic boundary conditions in the *y* direction. With these boundary conditions the IBM domain can be thought of as a cylinder, and could therefore represent a growing root or the developing intestine [22]. We use no-flux boundaries to augment the differences between GM1 and GM2 on the density of agents in apical growth [36, 37]. We only present results for linear and exponential domain growth in a two-species scenario, and in all simulations the domain grows to a horizontal length of 1500 lattice sites in the *x*-axis before the simulation is terminated. Both agent species are, on average, initially placed uniformly at random at densities of 0.05 (giving a total initial agent density of 0.1). All figures presented in this section are column averages taken from 1000 IBM repeats.

In Fig. 9 (a) and (b) density profiles for exponential enteric growth are shown. When GM1 is implemented species *A* dominates across the domain, although the density of species *A* is reduced in the region of high growth (see Fig. 9 (a)). However, in Fig. 9 (b) GM2 is implemented and causes species *B* to have a higher density in the middle region of the domain. In Fig. 9 (c) and (d) the density profiles for linear enteric growth are shown. As before, when GM1 is implemented this enables species *A* to dominate across the domain (see Fig. 9 (c)). However, in Fig. 9 (d) GM2 is implemented and this causes species *B* to have a slightly higher density in the middle region of the domain. Figures for apical non-uniform domain growth can be found in the supplementary information (Section S5).

**Figure 9:**
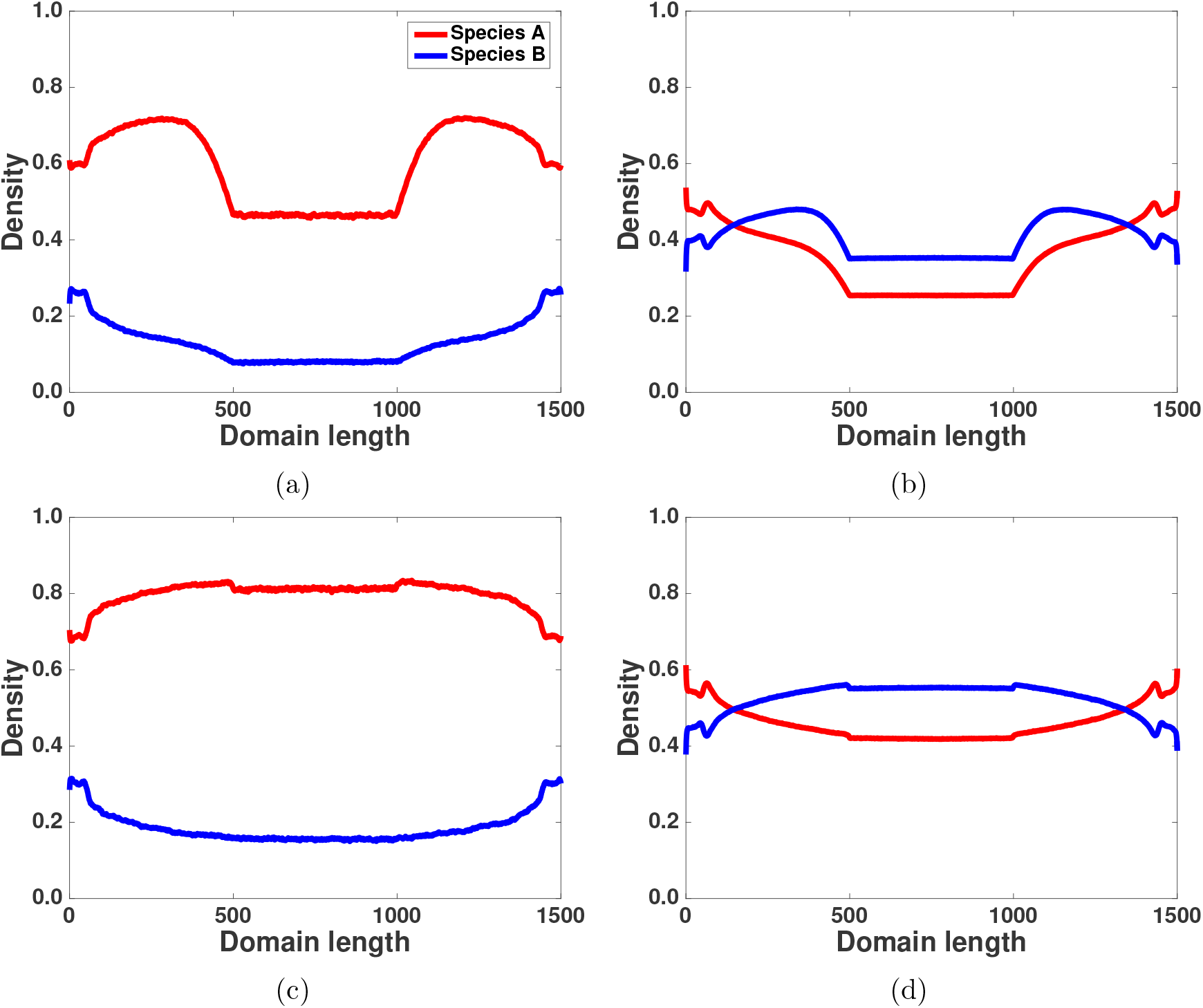
(Colour online): Enteric domain growth. (a) GM1 exponential domain growth, (b) GM2 exponential domain growth, (c) GM1 linear domain growth, (d) GM2 linear domain growth. The parameters for all panels are 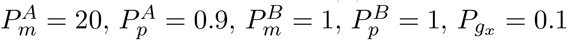.

In conclusion, with initial conditions that are, on average, spatially uniform for two species non-uniform growth can establish spatial variation in species density across the domain given certain parameter values. The development of this spatial variability is directly attributable to the differing effects of GM1 and GM2 on spatial correlations between agents in the IBM, as evidenced by the fact that when implemented with the same parameter values GM1 and GM2 produce different agent density profiles.

## 4 Discussion and conclusion

In this work we have studied the effect of two growth mechanisms on spatial correlations in agent populations containing multiple species. We chose two different, yet potentially biologically relevant growth mechanisms [23–25], to highlight how understanding the form of the domain growth in biological systems is important. Biologically, our growth mechanisms are simple descriptions of growth in the underlying tissue upon or within which a cell population is situated, a scenario often associated with migratory cell populations such as neural crest stem cells during embryonic development [5, 8, 15]. It is important to acknowledge that in reality it is unlikely that domain growth in biological systems is captured by algorithms as simple as GM1 and GM2. However, more realistic growth mechanisms may exist that exhibit similar effects on spatial correlations as GM1 and GM2.

Our key finding is that the specific type of growth mechanism can influence the dominant species, as shown in Figs. 4, 7 and 8. Under certain parameter regimes a more motile, slower proliferating species will dominate under growth mechanism GM1, whereas a less motile, faster proliferating species will dominate under growth mechanism GM2. This is because GM2 breaks down colinear correlations more effectively than GM1, and so benefits the faster proliferating species. Interestingly, this result suggests that the way in which a domain grows could play a role in determining cell population fates in biological systems, and to our knowledge is not a result that has been previously reported. To conclude our results section we studied some biologically motivated examples of non-uniform domain growth. We found that we were able to establish spatial variability in species densities (Fig. 9), and that this spatial variability changed depending on the way domain growth was implemented. This shows that non-uniform growth can establish spatially variable species densities on a domain, which is an intriguing result.

In this work all models studied are two-dimensional. The correlation ODE model has been derived for three dimensions on a non-growing domain [19], and so this is an obvious extension to the work presented here. In addition, we have also only considered the case when *P_gx_* = *P_gy_*. We did this to reduce the complexity of the equations, however, the results presented here could be extended to cases where *P_gx_* ≠ *P_c_*.

A final consideration is whether the work presented here could be extended to other types of model, such as an off-lattice IBM whereby agents can occupy any position in space (while taking volume exclusion into account, if necessary). Research has been directed towards including spatial structure in continuum approximations of off-lattice IBMs [38–40]. The effect of the domain growth mechanisms on the evolution of the agent density in an off-lattice IBM, as with the work presented here, will depend on how effectively the growth mechanisms break up spatial correlations established by agent proliferation.

## Acknowledgements

RJHR would like to thank the UK’s Engineering and Physical Sciences Research Council (EPSRC) for funding through a studentship at the Systems Biology programme of The University of Oxford’s Doctoral Training Centre. The authors would also like to thank Matthew Simpson and Deborah Markham for useful discussions.

## Supplementary information

### S1: Inclusion of agent proliferation, motility and death in the density functions

We first display how to include agent proliferation, motility and death in the individual density functions. To do so we require to introduce further notation. We indicate a site unoccupied by an agent in the following manner, 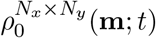, that is, the joint probability of lattice site **m** not being occupied on a domain of size *N_x_ × N_y_* at time *t*. We also introduce the summation, 
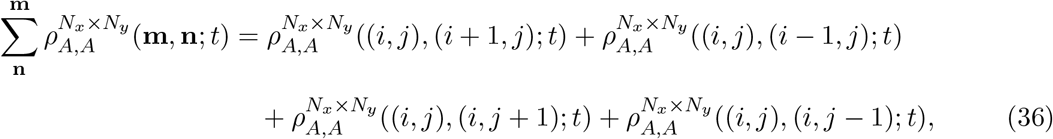
which indexes over the von Neumann neighbourhood of the site indicated as the upper index.

The sum of the individual density functions on a domain of size *N_x_ × N_y_* at [*t* + *δt*) for motile and proliferating agents is

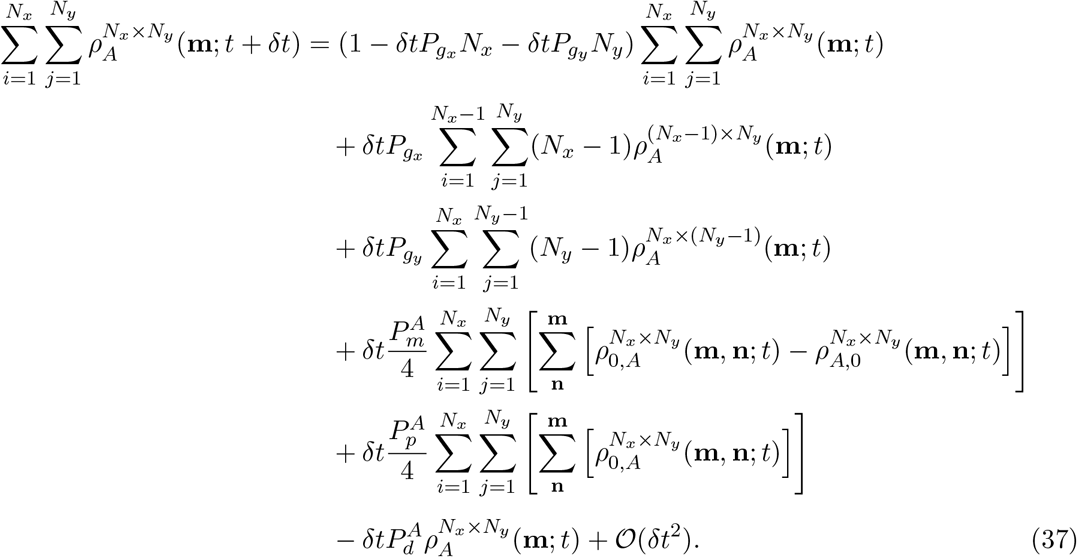

As we assume translational invariance Eq. (37) can be simplified to obtain

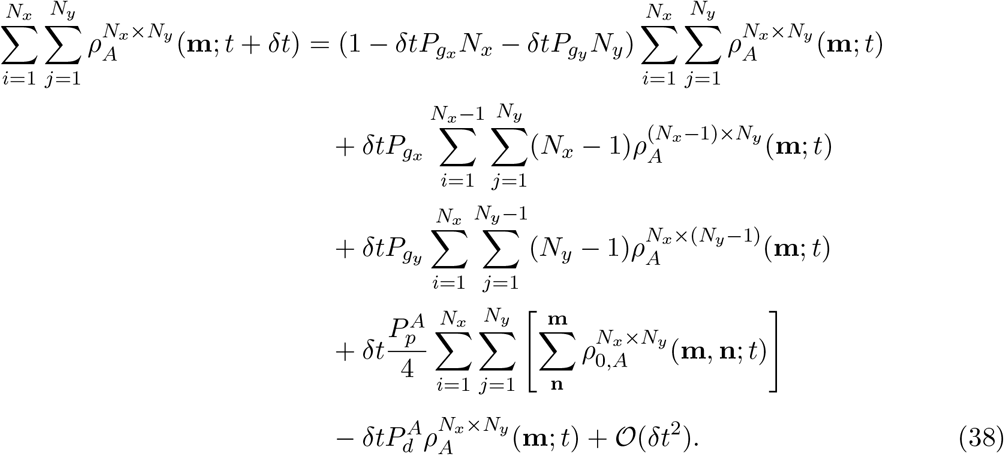

If we recognise that 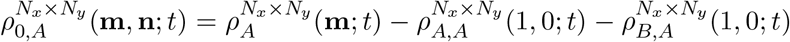 we can rewrite Eq. (38) as

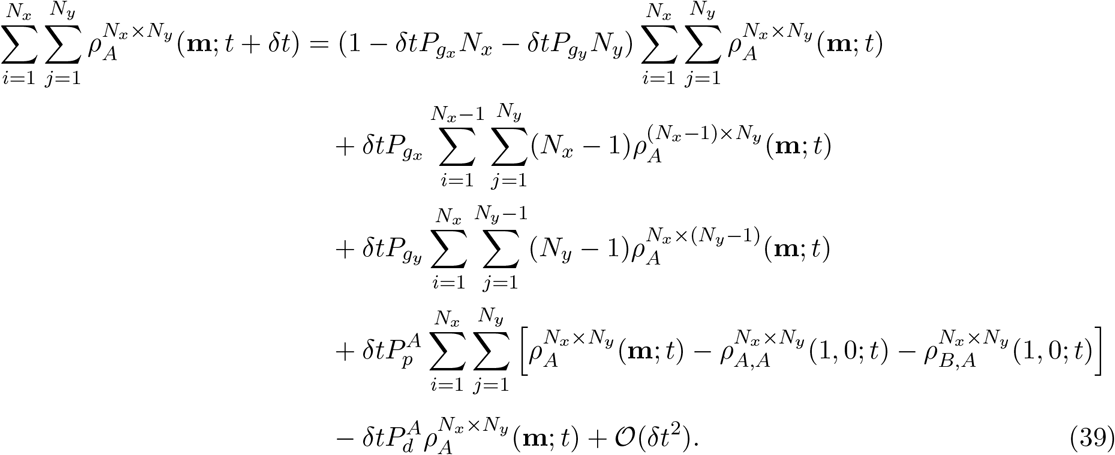

From Eq. (39) we can obtain

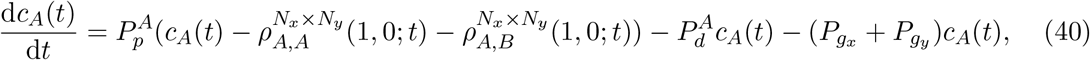
which is Eq. (27) in the text.

The derivation of the pairwise density functions for multispecies without domain growth can be found in [14]. However, we outline the derivation for the auto-correlations below. The addition of agent motility and proliferation is the same for GM1 and GM2. For agents colinear in the horizontal direction, that is, *r_y_* is zero, and *r_x_ >* 1, the evolution of the pairwise density functions for motile and proliferative agents with GM1 is

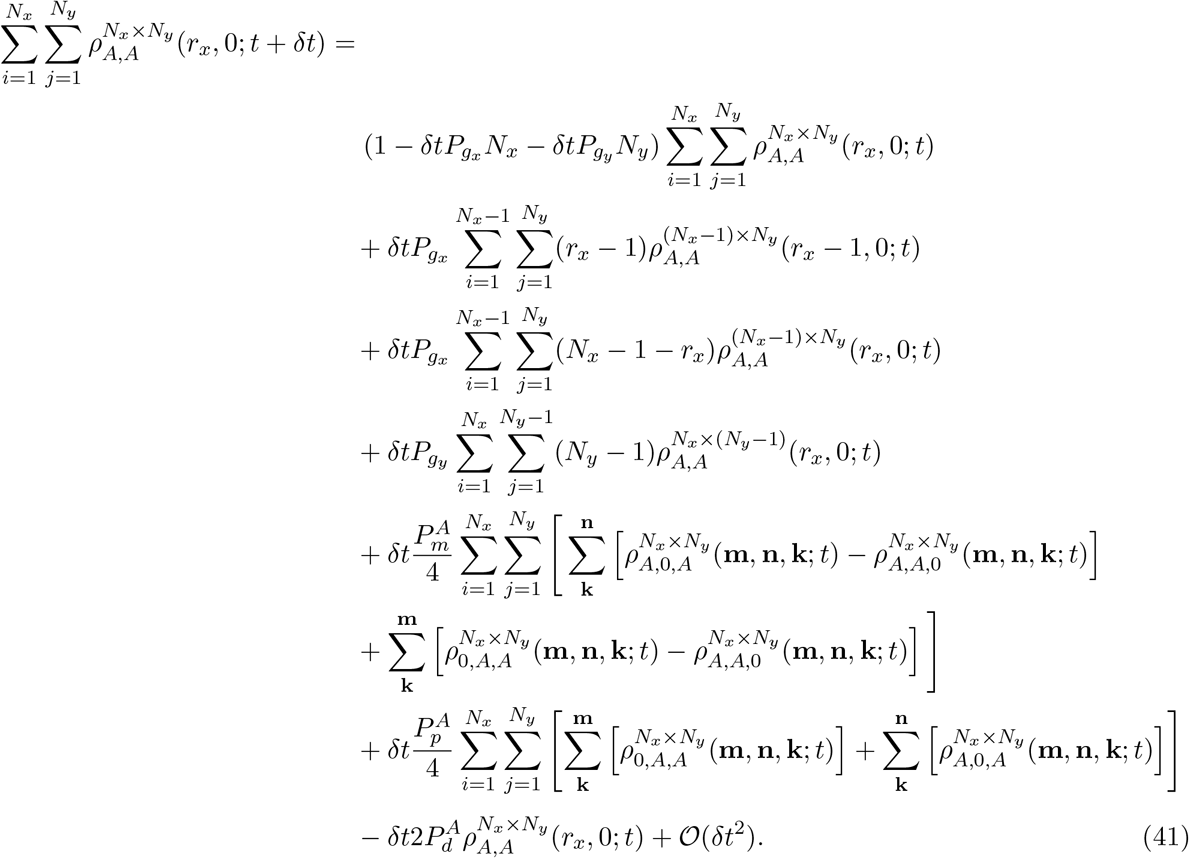

If *r_x_* = 1

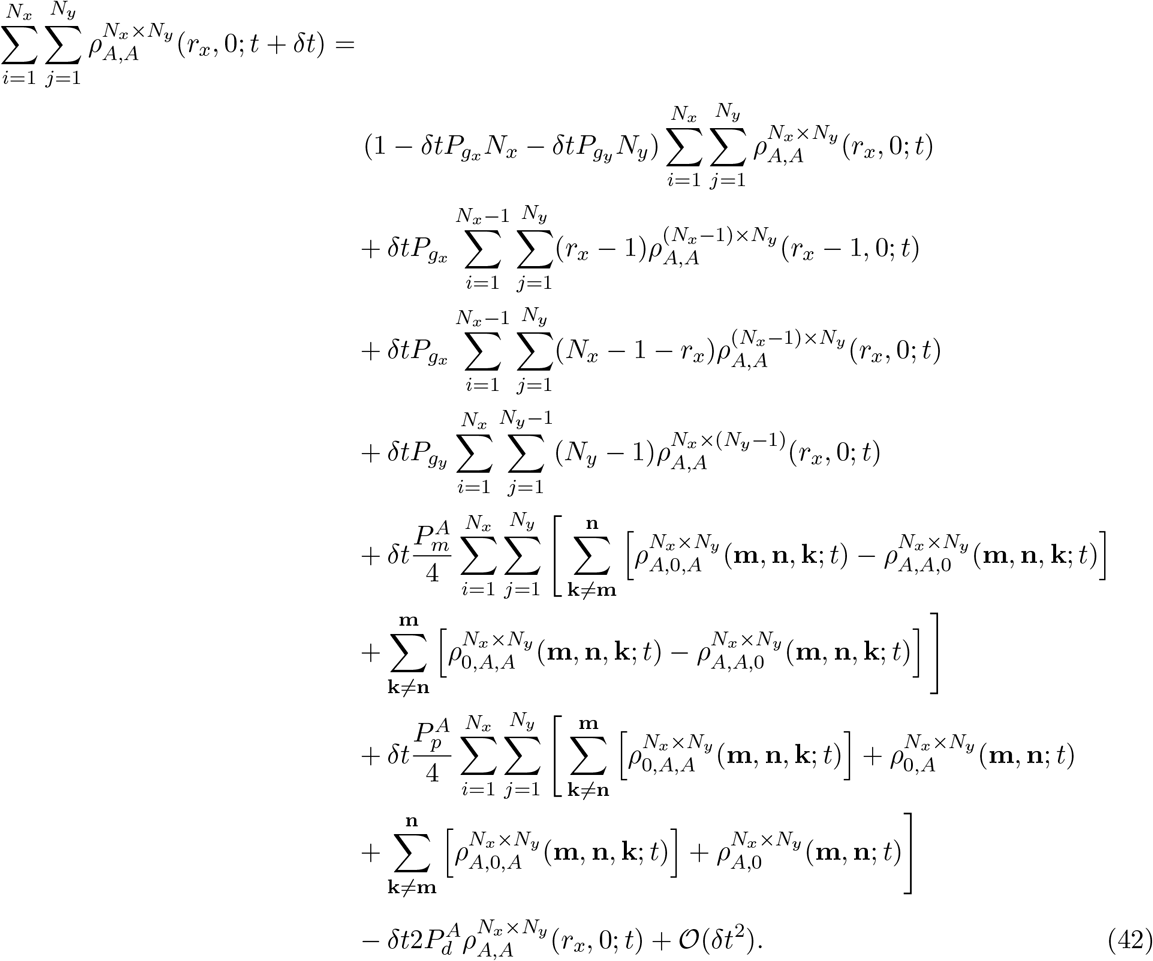
From which we can obtain the necessary equations [14].

### S2: Analysis of closure approximation

The approximation we employ in Eq. (6) in the main text has been previously published [16, 17, 26], and in Ross et al. [16] its accuracy for a one-dimensional lattice-based model has been demonstrated. In terms of one-point density functions this approximation 
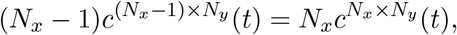
is a conservation statement that is exact in systems without proliferation. However, in the case of the pairwise density functions it is an approximation employed to make the equations tractable to solve numerically. Without this closure for the pairwise density functions the number of equations it is necessary to solve increases by a factor of (*n* + 1)*n/*2, where *n* is the total number of combinations of domain lengths that are simulated (computed).

For completeness we have attached further analysis of this closure approximation in the case of the individual density functions (i.e. *c^N_x_×N_y_^*(*t*)). We measure the relative error of the closure for the individual density functions

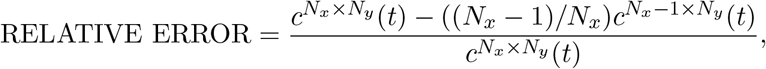
and plot the average relative error^7^ associated with each domain length for this closure in Fig. S1. It can be seen that the average relative error associated with the closure decreases as the domain grows larger, meaning the approximation improves as the domain grows. Other plots examining the error associated with the moment closures presented in this manuscript can be found in Ross et al. [16].

**Figure S1:**
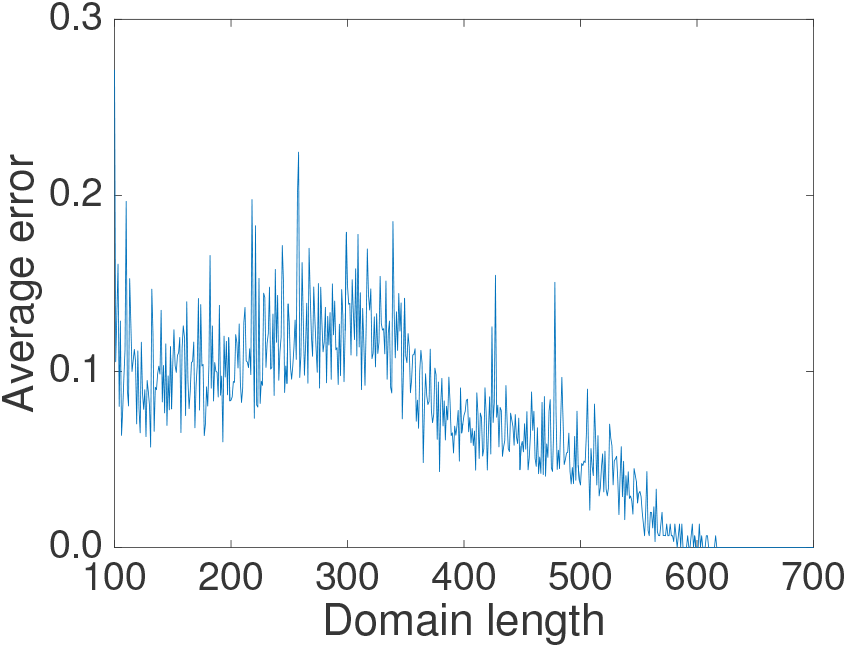
The average relative error associated with the domain length closure for individual density functions.

### S3: Growth mechanism 2

#### Diagonal component

It is simpler to derive the pairwise density functions for GM2 for a domain of size (*N_x_* + 1) *×* (*N_y_* + 1) (this avoids having (*N_x_ −* 1) in the denominator of a number of fractions). For GM2 it is also necessary to introduce some new notation. The distance of the lattice site **m** from the boundary (origin of growth) is *m_i_* and *m_j_*, i.e. the position of **m**, and the distance of the lattice site **n** from boundary, *n_i_* and *n_j_*, i.e. the position of **m** + (*r_x_, r_y_*). The evolution of the pairwise density functions for the diagonal component of GM2 is 
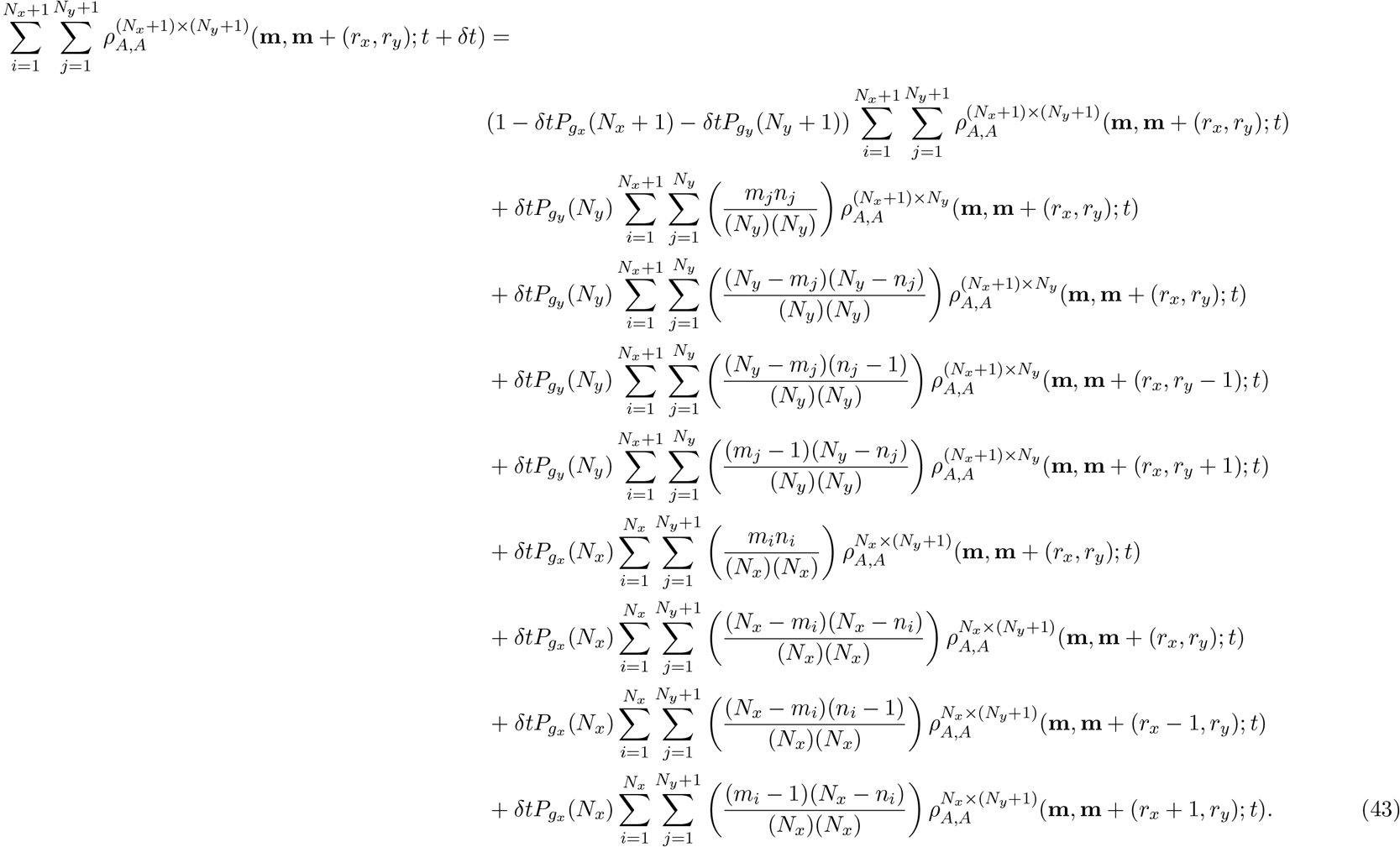

The terms on the RHS of Eq. (43) represent: i) no growth occurs in [*t* + *δt*), ii) the sum of pairwise density functions (*r_x_, r_y_*) apart on a domain of size (*N_x_* + 1)(*N_y_*) multiplied by the probability that a growth event occurs in the vertical direction and both agents are moved by it, iii) the sum of pairwise density functions (*r_x_, r_y_*) apart on a domain of size (*N_x_* + 1)(*N_y_*) multiplied by the probability that a growth event occurs in the vertical direction and both agents are not moved by it, iv) the sum of pairwise density functions (*r_x_, r_y_ −* 1) apart on a domain of size (*N_x_* + 1)(*N_y_*) multiplied by the probability that a growth event occurs in the vertical direction and **m**+(*r_x_, r_y_*) moves and **m** does not move (we can write this in terms of a displacement vector: (*r_x_, r_y_ −* 1)), and v) the sum of pairwise density functions (*r_x_, r_y_* + 1) apart on a domain of size (*N_x_* + 1)(*N_y_*) multiplied by the probability that a growth event occurs in the vertical direction and **m** moves and **m**+(*r_x_, r_y_*) does not move. The rest of the terms are the equivalent for growth in the horizontal direction. First, we assume translational invariance and simplify Eq. (43) to obtain 
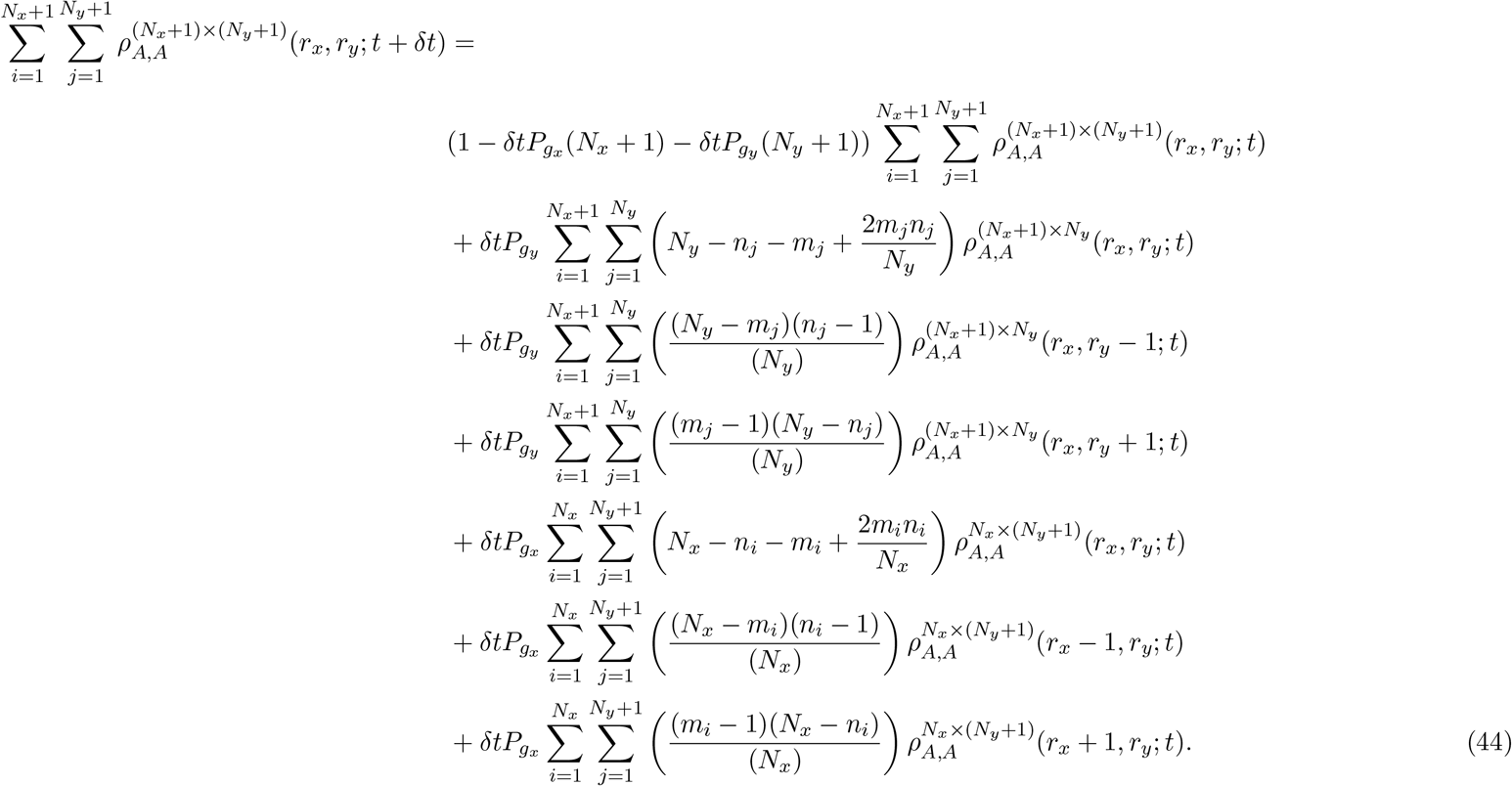

We rewrite Eq. (44) using 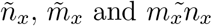, which are constants defined as 
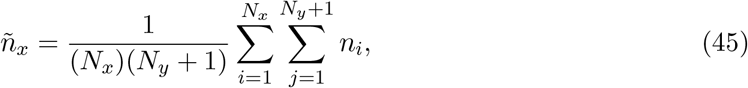
and 
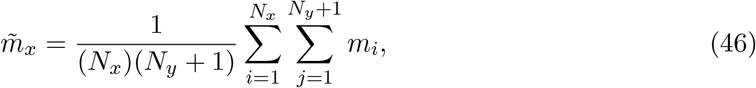
and 
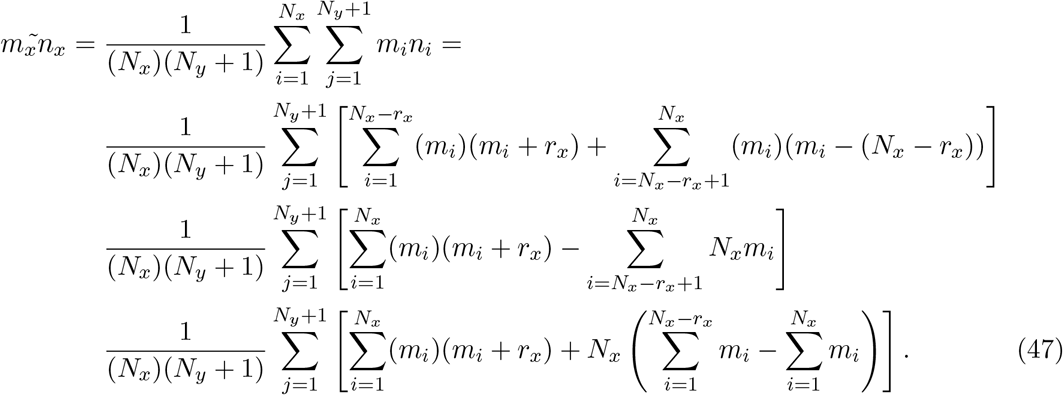
Eqs. (45)-(47) can be evaluated directly. If we do so we obtain: 
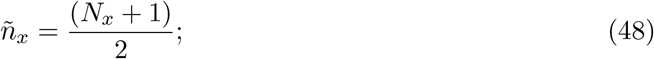

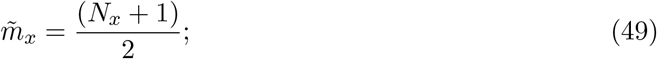
and 
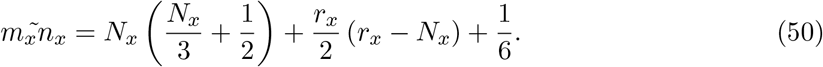

It is important to note that if in Eq. (50) we set *r_x_* = 0, for instance for colinear lattice sites, we obtain 
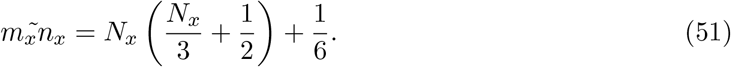

However, initially we substitute Eqs. (45)-(47) into Eq. (44) to obtain 
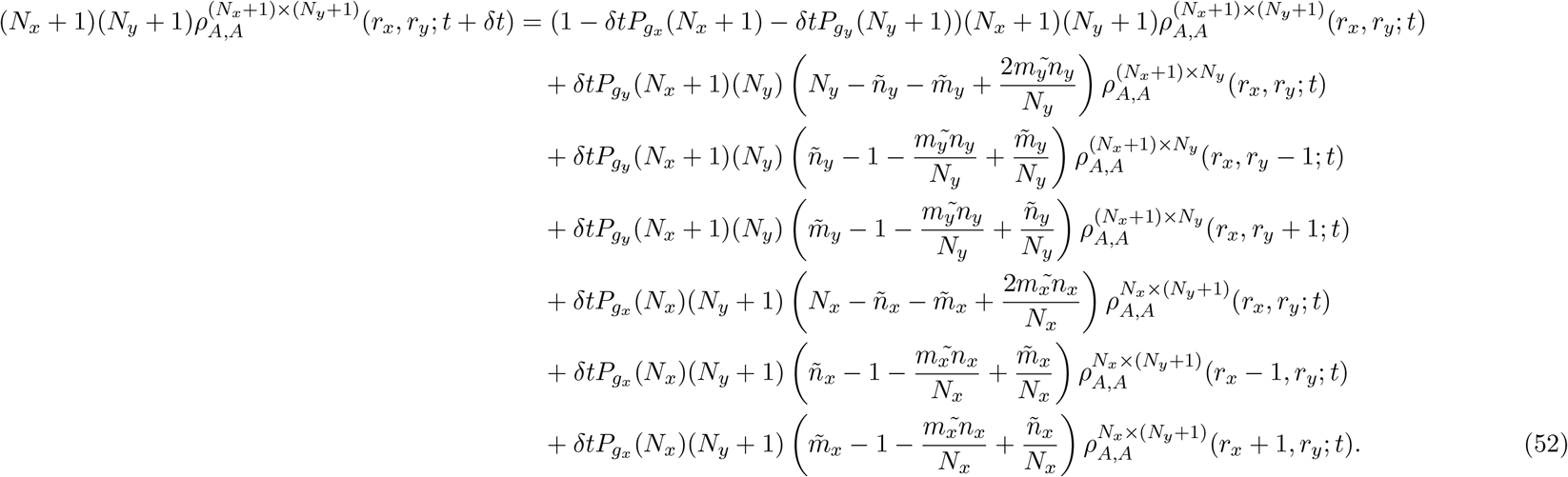

That Eqs. (44) and (52) are equivalent can be shown by substituting Eqs. (45)-(47) into Eq. (52).

If we now apply the approximation, 
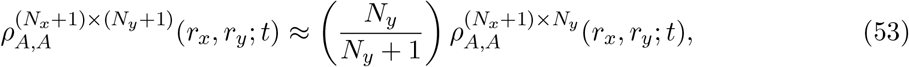
to Eq. (52) we obtain 
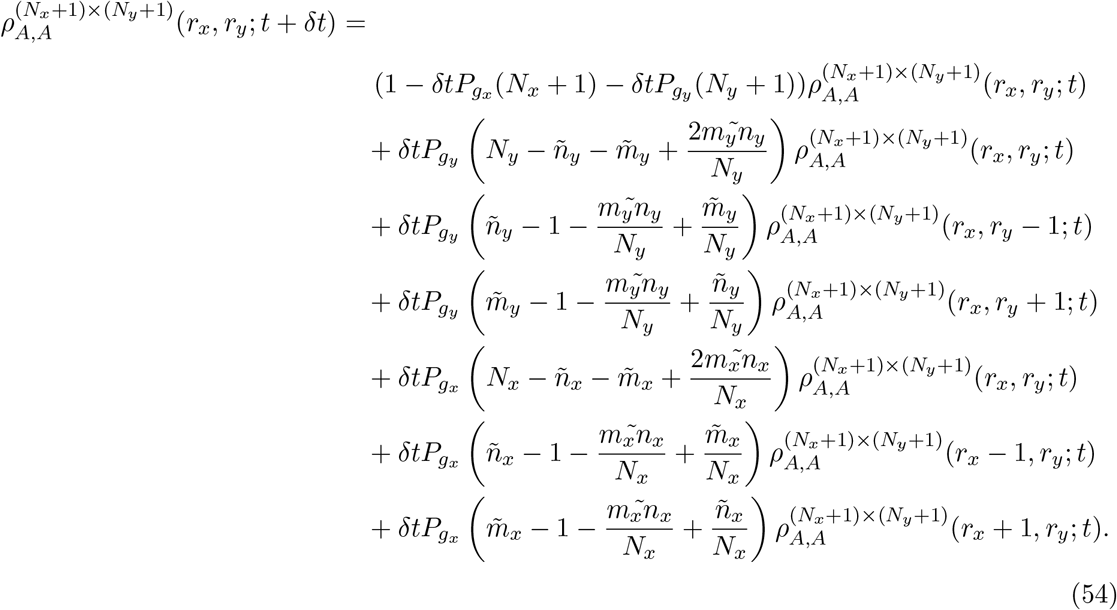

If we rearrange Eq. (54) and take the limit as *δt →* 0 we obtain 
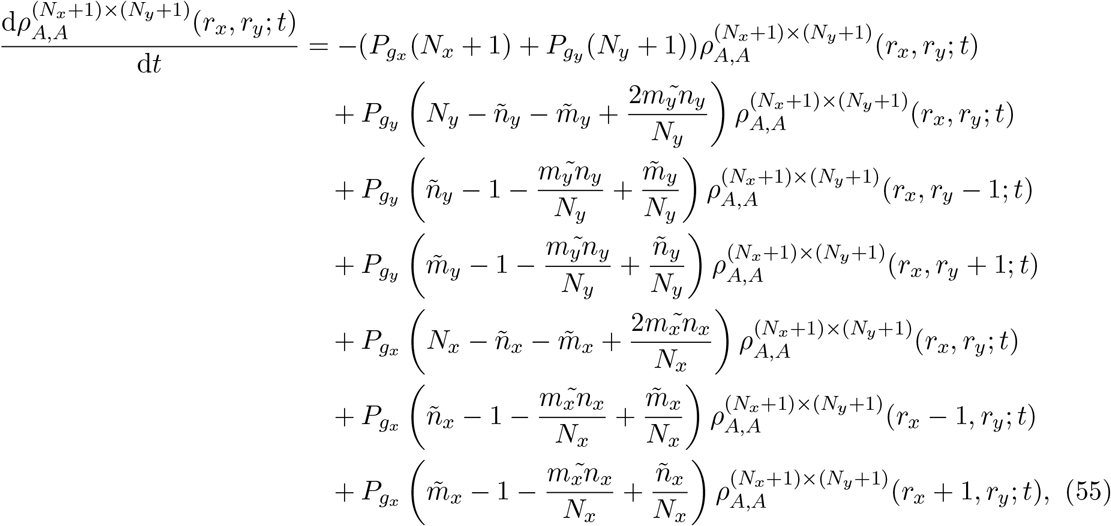
which we can simplify further to obtain 
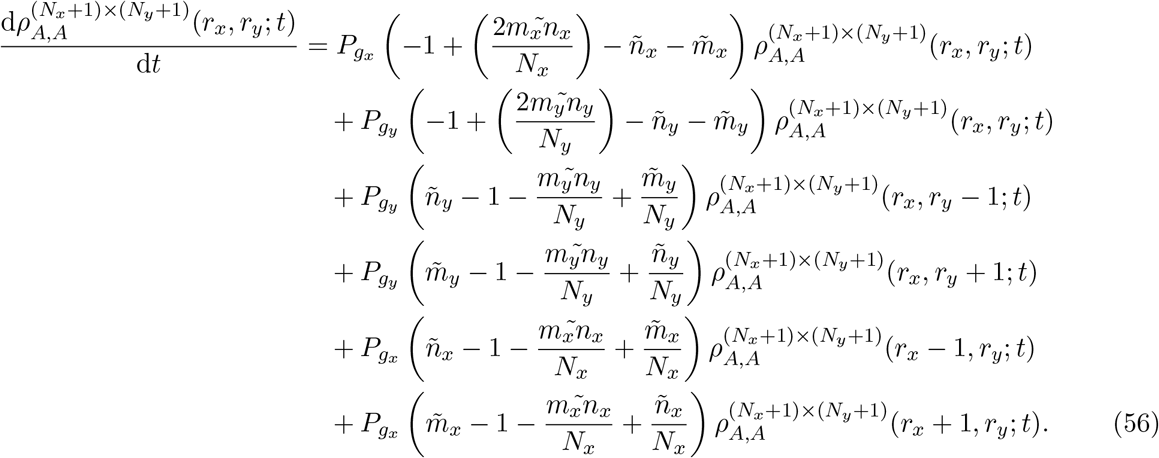

If we now evaluate Eqs. (45)-(47) with Eqs. (48)-(50) we obtain 
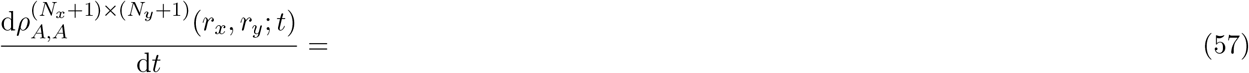

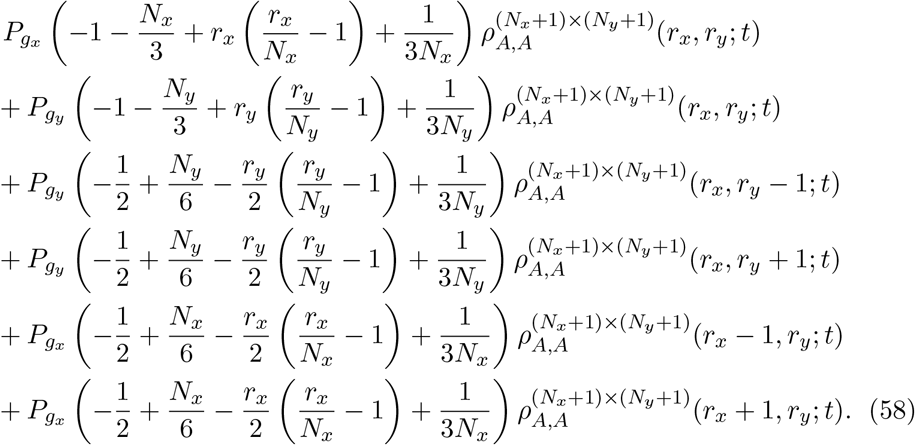

Equation (58) is Eq. (23) in the main text.

For the colinear component we have (assuming that lattice sites are horizontally colinear, i.e. *r_y_* = 0) 
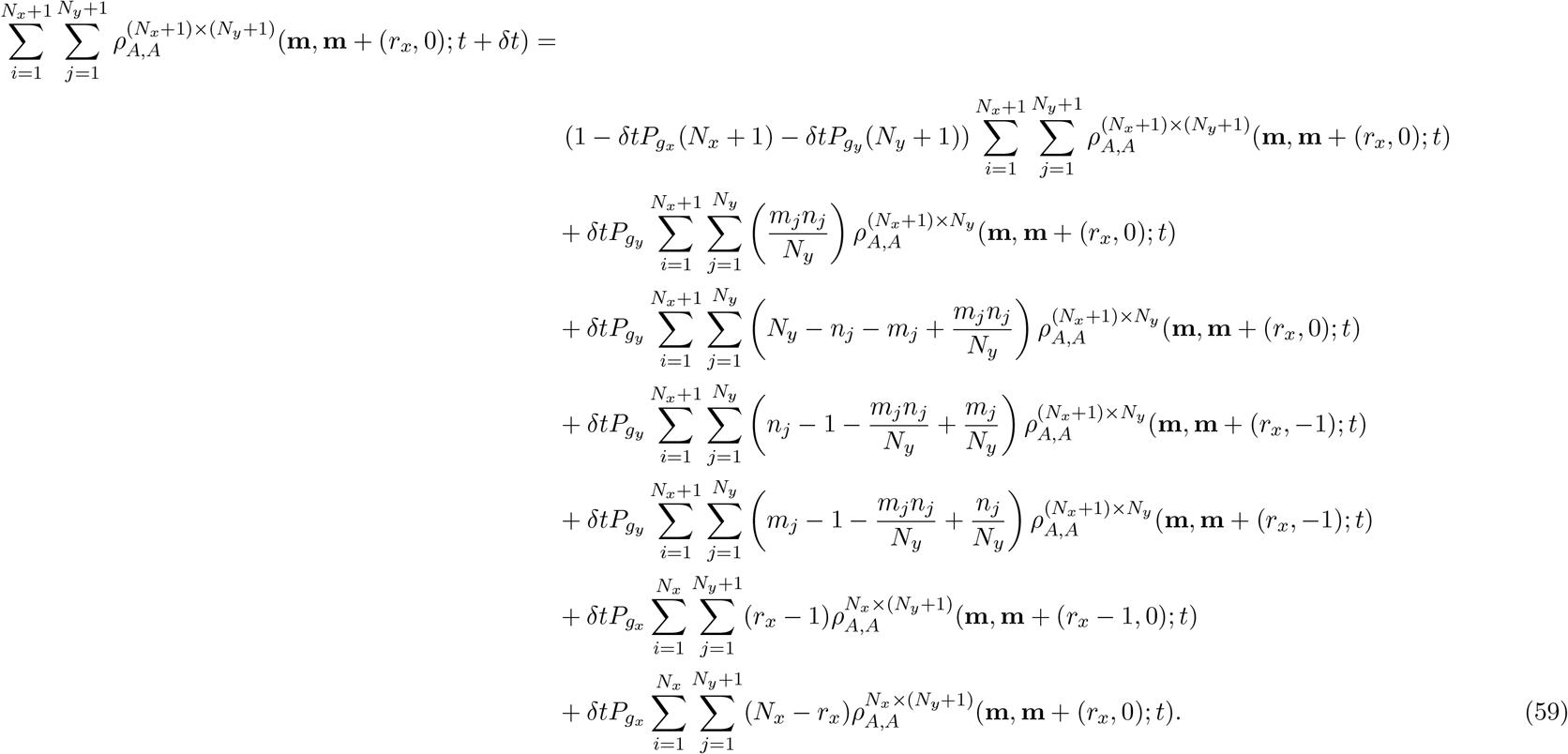

We rewrite Eq. (59) using Eqs. (45)-(47) to obtain 
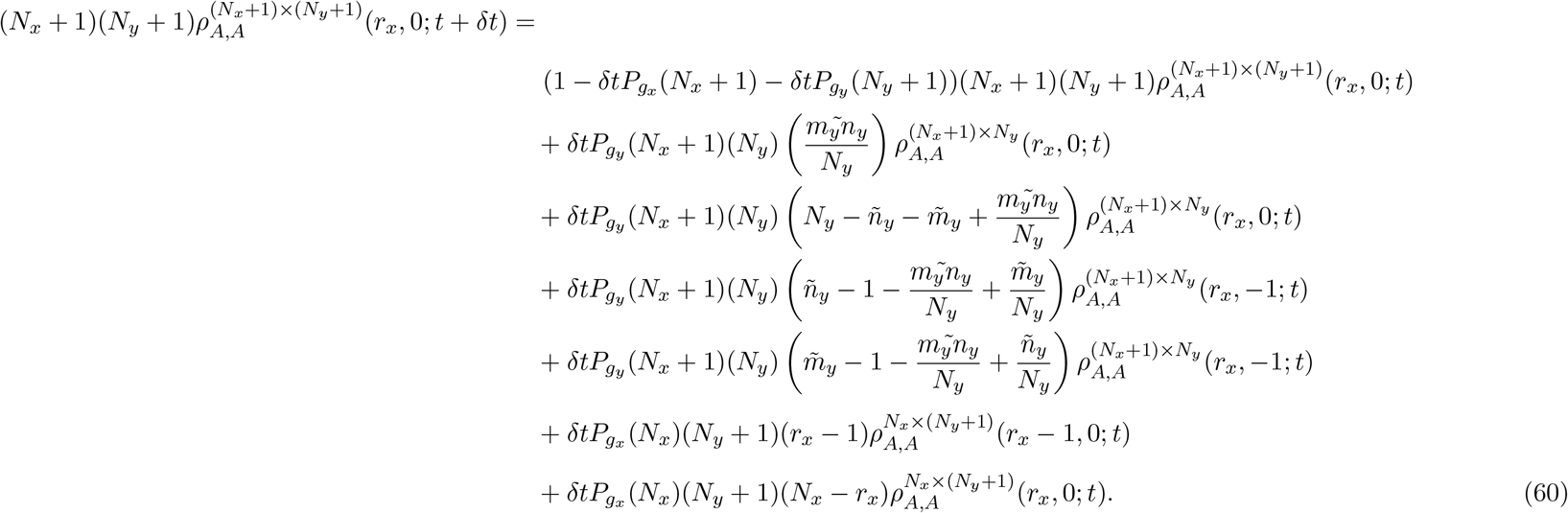

We now make the approximation Eq. (53) to obtain

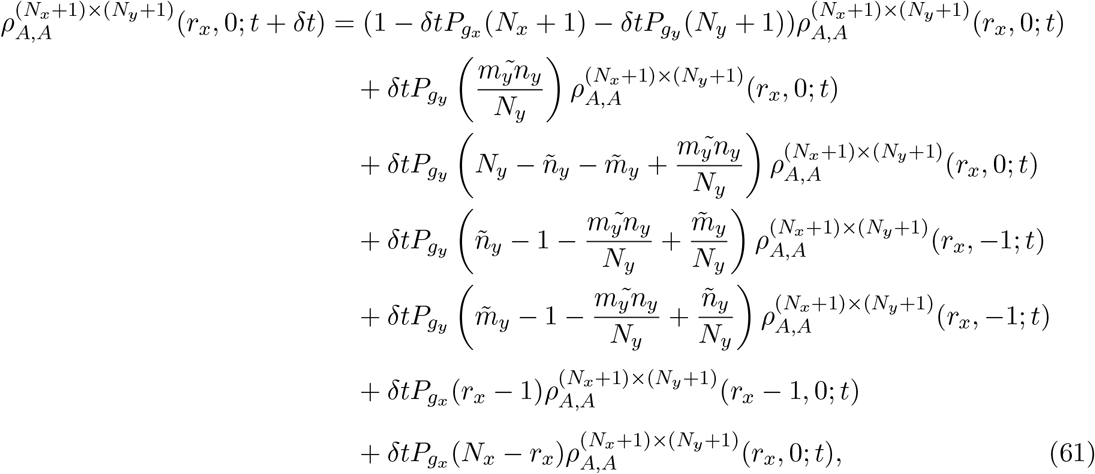
which we can simplify to obtain 
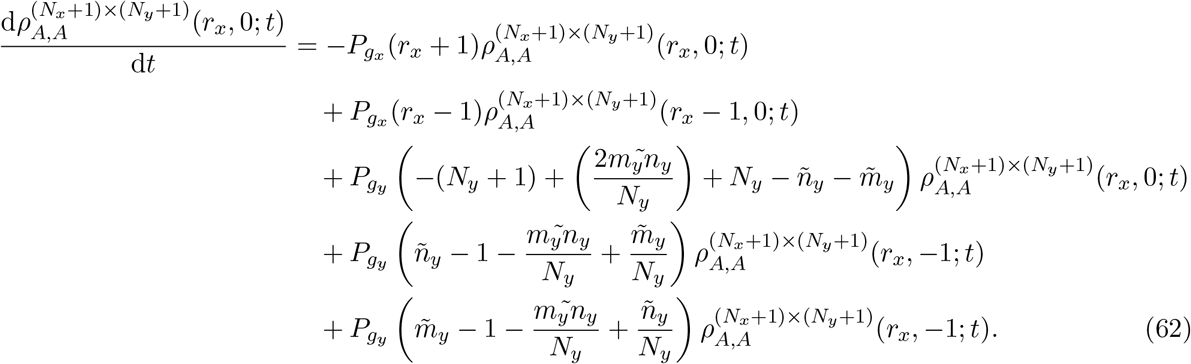

If we evaluate Eqs. (45)-(47) with (48)-(50) we obtain 
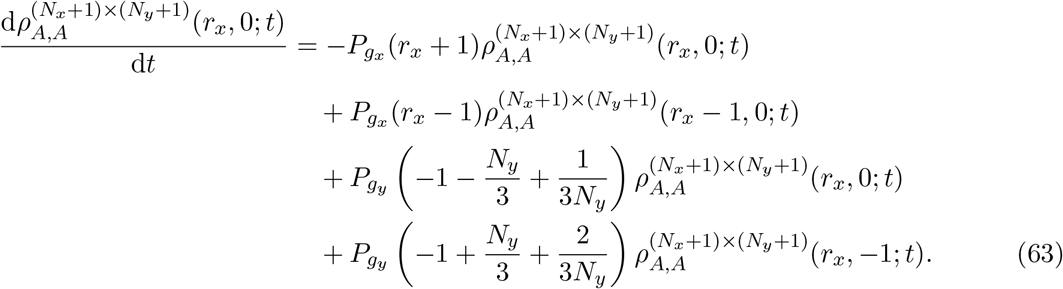
Equation (63) is Eq. (22) in the main text.

### S4: Parameter sweeps for GM1 and GM2

In Fig. S2 we present a parameter sweep for a domain of initial dimensions *N_x_* = 100 by *N_y_*= 100, 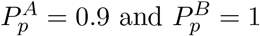, and sampled at simulation time *t* = 25 for GM1 and GM2. The coordinates of this parameter sweep are 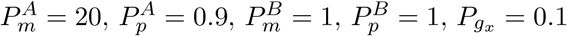 is plotted. Figure S2 shows that, depending on the growth mechanism implemented, species *A* or species *B* dominates the domain for certain parameter values at time *t* = 25. It is important to remember that for GM2 the dominant species at a given time point not only depends on both the motility and proliferation parameters associated with species *A* and *B*, but also the initial length of the domain (species densities under GM1 do not depend on the initial size of the domain, which Fig. 7 in the main text demonstrates). Therefore, Fig. S2 (b) would be different if the initial dimensions of the domain were altered. Both (a) and (b) in Fig. S2 would also be different if we sampled at an alternative simulation time.

**Figure S2:**
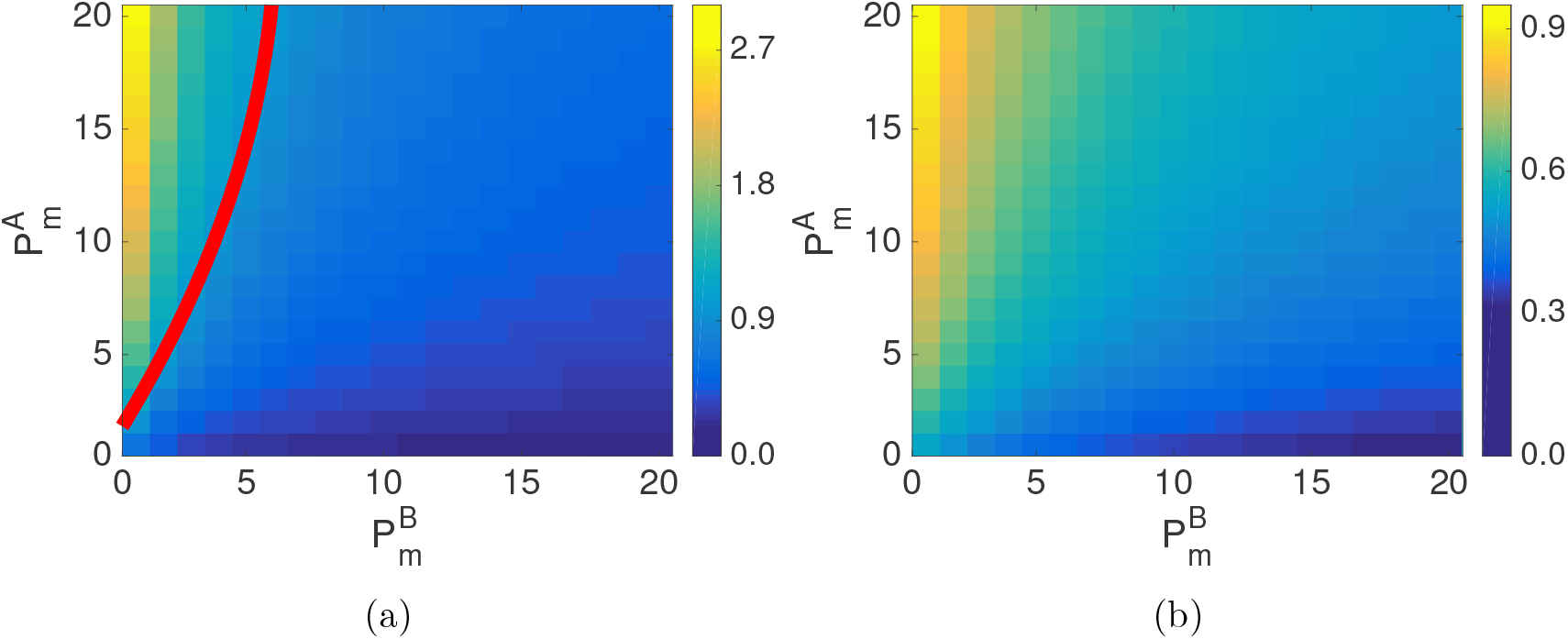
Parameter sweep for (a) GM1 and (b) GM2. *P_gx_* = 0.1 and *P_gy_* = 0.1 and domain growth is exponential. 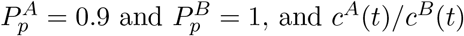 is plotted. The red contour line in (a) indicates 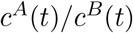 = 1.

### S5: Non-uniform domain growth: apical

In Fig. S3 (a) and (b) density profiles for exponential apical growth are shown. In Fig. S3 (a) GM1 is implemented and this enables species *A* to dominate across the domain. However in Fig. S3 (b) GM2 is implemented and this enables species *A* and *B* to have similar densities in the middle region of the domain. Similarly, in Fig. S3 (c) and (d), density profiles for linear apical growth are shown. In Fig. S3 (c) GM1 is implemented and, much like exponential growth, this results in species *A* to dominate across the domain. In contrast, in Fig. S3 (d) GM2 is implemented and this results in species *A* and *B* to have similar densities in the middle region of the growing domain. This result suggests that the form of growth in apical growth can determine the dominant species at the interface of two differently growing regions, and could have interesting implications in biological systems with apical growth [36, 37].

**Figure S3:**
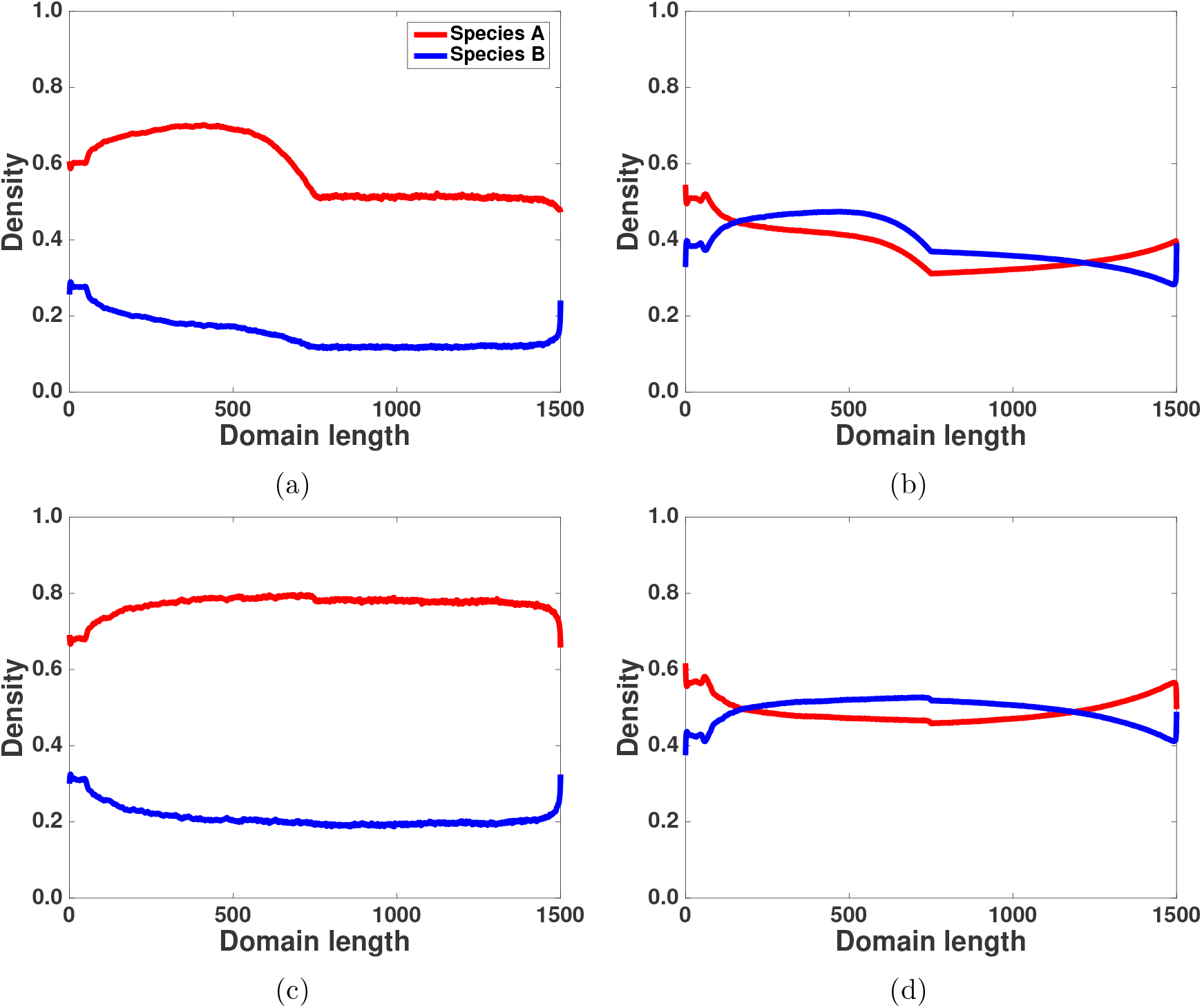
(Colour online): Apical domain growth. (a) GM1 exponential domain growth, (b) GM2 exponential domain growth, (c) GM1 linear domain growth, (d) GM2 linear domain growth. The parameters for all panels are 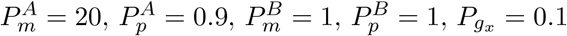 and *P*_gy_ = 0.1.

## Footnotes

1 We present the details of how to include the effects of agent motility, proliferation, and death, in the density functions in the supplementary information (Section S1).

2 As 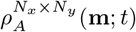 denotes a probability its domain of definition is [0,1].

3 A simpler (one-dimensional) version of this derivation can be found in Ross et al. [16

4 We assume translational invariance throughout this work because the initial agent density for all simulations in the IBM is achieved by populating lattice sites uniformly at random until the required density is achieved.

5 These approximations sensibly imply that domain growth ‘dilutes’ pairwise agent densities.

6 The agent motility and proliferation parameters we implement in this section have been chosen to illustrate a given effect. In Section S4 of the supplementary information we demonstrate that the effects we present in this manuscript can be reproduced by many other parameter values.

7 To compute the average relative error we proceed as follows: we initialise a domain of length *N_x_* = 100, *N*_y_ = 100, with *P_gx_* = 0.1 and *P_gy_* = 0 (i.e. no growth in the *y* direction), 500 initial agents (assigned uniformly at random to lattice sites) and *P_p_* = 1 and *P_m_* = 1. We calculate the relative error associated with the closure for all domain sizes, for example *c*^109×100^(*t*) = (110/109)*c*^110×100^(*t*), for the duration of the simulation sampling at equally spaced time intervals. We then sum the absolute values of the relative error for each domain length closure for each time point, and divide by the number of samples (time points) to generate the average relative error associated with this closure over the time course of the simulation.

## References

[1] R. Satija and A. K. Shalek. Heterogeneity in immune responses: from populations to single cells. Trends in Immunology, 35(5):219–229, 2014.

[2] M. F. Bear, B. W. Connors, and M. A. Paradiso. Neuroscience: exploring the brain. Lippincott Williams and Wilkins, Philadelphia, 3rd edition, 2007.

[3] G. Schram, M. Pourrier, P. Melnyk, and S. Nattel. Differential distribution of cardiac ion channel expression as a basis for regional specialization in electrical function. Circulation Research, 90(9):939–950, 2002.

[4] A. Marusyk, V. Almendro, and K. Polyak. Intra-tumour heterogeneity: a looking glass for cancer? Nature Reviews Cancer, 12(5):323–334, 2012.

[5] R. McLennan, L. Dyson, K. W. Prather, J. A. Morrison, R. E. Baker, P. K. Maini, and P. M. Kulesa. Multiscale mechanisms of cell migration during development: theory and experiment. Development, 139(16):2935–2944, 2012.

[6] R. McLennan, L. J. Schumacher, J. A. Morrison, J. M. Teddy, D. A. Ridenour, A. C. Box, C. L. Semerad, H. Li, W. McDowell, D. Kay, P. K. Maini, R. E. Baker, and P. M. Kulesa. Neural crest migration is driven by a few trailblazer cells with a unique molecular signature narrowly confined to the invasive front. Development, 142(11):2014–2025, 2015.

[7] R. McLennan, L. J. Schumacher, J. A. Morrison, J. M. Teddy, D. A. Ridenour, A. C. Box, C. L. Semerad, H. Li, W. McDowell, D. Kay, P. K. Maini, R. E. Baker, and P. M. Kulesa. VEGF signals induce trailblazer cell identity that drives neural crest migration. Developmental Biology, 407(1):12–25, 2015.

[8] R. L. Mort, R. J. H. Ross, K. J. Hainey, O. Harrison, M. A. Keighren, G. Landini, R. E. Baker, K. J. Painter, I. J. Jackson, and C. A. Yates. Reconciling diverse mammalian pigmentation patterns with a fundamental mathematical model. Nature Communications, 7(10288), 2016.

[9] B. Waclaw, I. Bozic, M. E. Pittman, R. H. Hruban, B. Vogelstein, and M. A. Nowak. A spatial model predicts that dispersal and cell turnover limit intratumour heterogeneity. Nature, 525(7568):261–264, 2015.

[10] D. J. G. Agnew, J. E. F. Green, T. M. Brown, M. J. Simpson, and B. J. Binder. Distinguishing between mechanisms of cell aggregation using pair-correlation functions. Journal of Theoretical Biology, 352:16–23, 2014.

[11] B. J. Binder and M. J. Simpson. Quantifying spatial structure in experimental observations and agent-based simulations using pair-correlation functions. Physical Review E, 88(2): 022705, 2013.

[12] B. J. Binder and M. J. Simpson. Spectral analysis of pair-correlation bandwidth: application to cell biology images. Royal Society Open Science, 2(2):140494, 2015.

[13] R. J. H. Ross, C. A. Yates, and R. E. Baker. Inference of cell–cell interactions from population density characteristics and cell trajectories on static and growing domains. Mathematical Biosciences, 264:108–118, 2015.

[14] D. C. Markham, M. J. Simpson, P. K. Maini, E. A. Gaffney, and R. E. Baker. Incorporating spatial correlations into multispecies mean-field models. Physical Review E, 88(5):052713, 2013.

[15] B. J. Binder, K. A. Landman, D. F. Newgreen, J. E. Simkin, Y. Takahashi, and D. Zhang. Spatial analysis of multi-species exclusion processes: application to neural crest cell migration in the embryonic gut. Bulletin of Mathematical Biology, 74(2):474–90, 2012.

[16] R. J. H. Ross, C. A. Yates, and R. E. Baker. The effect of domain growth on spatial correlations. Physica A, 466:334–345, 2016.

[17] R. J. H. Ross, R. E. Baker, and C.A. Yates. How domain growth is implemented determines the long term behaviour of a cell population through its effect on spatial correlations. Physical Review E, 94(1):012408, 2016.

[18] R. E. Baker and M. J. Simpson. Correcting mean-field approximations for birth-death-movement processes. Physical Review E, 82(4):041905, 2010.

[19] D. C. Markham, M. J. Simpson, and R. E. Baker. Simplified method for including spatial correlations in mean-field approximations. Physical Review E, 87(6):062702, 2013.

[20] T. M. Liggett. Stochastic Interacting Systems: Contact, Voter, and Exclusion Processes. Springer-Verlag, Berlin, 1999.

[21] D. T. Gillespie. Exact stochastic simulation of coupled chemical reactions. Journal of Physical Chemistry, 81(25):2340–2361, 1977.

[22] B. J. Binder, K. A. Landman, M. J. Simpson, M. Mariani, and D. F. Newgreen. Modeling proliferative tissue growth: A general approach and an avian case study. Physical Review E, 78(3):031912, 2008.

[23] B. J. Thomas, D. A. Gunning, J. Cho, and S. L. Zipursky. Cell cycle progression in the developing Drosophila eye: roughex encodes a novel protein required for the establishment of G1. Cell, 77(7):1003–1014, 1994.

[24] J. P. Kumar. My what big eyes you have: How the Drosophila retina grows. Developmental Neurobiology, 71(12):1133–1152, 2011.

[25] C. J. Barelle, E. A. Bohula, S. J. Kron, D. Wessels, D. R. Soll, A. Schäfer, A. J. P. Brown, and N. A. R. Gow. Asynchronous cell cycle and asymmetric vacuolar inheritance in true hyphae of Candida albicans. Eukaryotic cell, 2(3):398–410, 2003.

[26] R. E. Baker, C. A. Yates, and R. Erban. From microscopic to macroscopic descriptions of cell migration on growing domains. Bulletin of Mathematical Biology, 72(3):719–762, 2010.

[27] M. A. M. de Aguiar, E. M. Rauch, and Y. Bar-Yam. Invasion and extinction in the mean-field approximation for a spatial host-pathogen model. Journal of Statistical Physics, 114(5–6):1417–1451, 2004.

[28] D. Hiebeler. Stochastic spatial models: From simulations to mean-field and local structure approximations. Journal of Theoretical Biology, 187(3):307–319, 1997.

[29] H. Fuks and A. T. Lawniczak. Individual-based lattice model for spatial spread of epidemics. Discrete Dynamics in Nature and Society, 6(3):191–200, 2001.

[30] J. Mai, V. N. Kuzovkov, and W. von. Niessen. A theoretical stochastic model for the A+1/2B → 0 reaction. Journal of Chemical Physics, 98(12):10017, 1993.

[31] J. Mai, V. N. Kuzovkov, and W. von. Niessen. A general stochastic model for the description of surface reaction systems. Physica A, 203:298–315, 1994.

[32] M. J. Simpson and R. E. Baker. Corrected mean-field models for spatially dependent advection-diffusion-reaction phenomena. Physical Review E, 83(5):051922, 2011.

[33] S. T. Johnston, M. J. Simpson, and R. E. Baker. Mean-field descriptions of collective migration with strong adhesion. Physical Review E, 85(5):051922, 2012.

[34] M. J. Simpson, A. Merrifield, K. A. Landman, and B. D. Hughes. Simulating invasion with cellular automata: Connecting cell-scale and population-scale properties. Physical Review E, 76:021918, 2007.

[35] K. A. Landman and A. E. Fernando. Myopic random walkers and exclusion processes: Single and multispecies. Physica A, 390(21-22):3742–3753, 2011.

[36] S. Perilli, R. Di Mambro, and S. Sabatini. Growth and development of the root apical meristem. Current Opinion in Plant Biology, 15(1):17–23, 2012.

[37] J. F. Fallon, A. Lopez, M. A. Ros, M. P. Savage, B. B. Olwin, and B. K. Simandl. FGF2: apical ectodermal ridge growth signal for chick limb development. Science, 264(5155): 104–107, 1994.

[38] R. Law, D. J. Murrell, and U. Dieckmann. Population growth in space and time: spatial logistic equations. Ecology, 84(1):252–262, 2003.

[39] D. J. Murrell. Local spatial structure and predator-prey dynamics: counterintuitive effects of prey enrichment. The American Naturalist, 166(3):354–367, 2005.

[40] D. J. Murrell. When does local spatial structure hinder competitive coexistence and reverse competitive hierarchies? Ecology, 91(6):1605–1616, 2010.

